# Regulation of expression site and generalizability of experience-dependent plasticity in visual cortex

**DOI:** 10.1101/773275

**Authors:** Crystal L. Lantz, Sachiko Murase, Elizabeth M. Quinlan

## Abstract

The experience-dependent decrease in stimulus detection thresholds that underly perceptual learning can be induced by repetitive exposure to a visual stimulus. Robust stimulus-selective potentiation of visual responses is induced in the primary mouse visual cortex by repetitive low frequency visual stimulation (LFVS). How the parameters of the repetitive visual stimulus impact the site and specificity of this experience-dependent plasticity is currently a subject of debate. Here we demonstrate that the stimulus selective response potentiation induced by repetitive low frequency (1 Hz) stimulation, which is typically limited to layer 4, shifts to superficial layers following manipulations that enhance plasticity in primary visual cortex. In contrast, repetitive high frequency (10 Hz) visual stimulation induces response potentiation that is expressed in layers 4 and 5/6, and generalizes to novel visual stimuli. Repetitive visual stimulation also induces changes in the magnitude and distribution of oscillatory activity in primary visual cortex, however changes in oscillatory power do not predict the locus or specificity of response potentiation. Instead we find that robust response potentiation is induced by visual stimulation that resets the phase of ongoing gamma oscillations. Furthermore, high frequency, but not low frequency, repetitive visual stimulation entrains oscillatory rhythms with enhanced sensitivity to phase reset, such that familiar and novel visual stimuli induce similar visual response potentiation.

## Introduction

Repeated exposure to a sensory stimulus induces perceptual learning, characterized by a long-term decrease in detection or discrimination thresholds for the experienced stimulus (Karni and Sagi, 1993; Poggio et al., 1992; Sagi, 2011; Schoups et al., 1995; Seitz, 2017). Perceptual learning has been demonstrated in all sensory modalities and does not decline with age (Brickwedde et al., 2019; Karawani et al., 2016). In animal models, stimulus-selective perceptual learning is characterized as a robust enhancement of sensory responses following active sensory discrimination or passive stimulus exposure (Crist et al., 2001; Frenkel et al., 2006; Ren et al., 2016). Accordingly, perceptual learning has the potential to be administered therapeutically in an attempt to improve sensory discrimination in disorders such as amblyopia or age-related sensory loss. However, the rigid stimulus-selectivity of perceptual learning constrains clinical utility (Deveau and Seitz, 2014; Karawani et al., 2016).

Work in animal models is beginning to lend insight into the mechanism by which repetitive stimulation induces stimulus-selective response enhancement. In mice, repetitive presentation of high contrast square wave gratings reversing at low frequency (LFVS; 1 Hz reversal /2 Hz screen refresh; 100s of presentations/day for up to 5 days) induces robust and stimulus-selective enhancement of visual responses (Andermann et al., 2010; Aton et al., 2014; Frenkel et al., 2006). This stimulus-selective response potentiation (SRP) has been demonstrated physiologically as an increase in the amplitude of visually evoked potentials (VEPs) in layer 4, an increase in the firing rate of regular spiking (RS) neurons throughout primary visual cortex and an increase in orientation tuning of regular-spiking neurons tuned to the familiar stimulus (Cooke et al., 2015; Frenkel et al., 2006). Importantly, LFVS also induces a decrease in the magnitude of the behavioral ‘vidget,’ which reports stimulus-selective visual response habituation (Cooke et al., 2015). SRP emerges slowly over days and requires coherent firing between cortico-thalamic neurons in layer 6 and neurons in the LGN during sleep for consolidation (Aton et al., 2014; Durkin et al., 2017). Importantly, SRP is robust in adults, indicating a distinction from other forms of synaptic plasticity in primary visual cortex that are constrained by age (Daw et al., 1992; Espinosa and Stryker, 2012; Gordon and Stryker, 1996; Jiang et al., 2007). SRP is also highly selective for the characteristics of the familiar visual stimulus such as orientation, contrast, or spatial frequency. Indeed, rotation of an oriented grating as little as 5 degrees is sufficient to block the expression of SRP (Cooke and Bear, 2010). The selectivity of SRP replicates many forms of visual perceptual learning in humans, which has been shown to improve discrimination for characteristics of the familiar stimulus including orientation, texture contrast and spatial frequency (Fiorentini and Berardi, 1981; Karni et al., 1994; Poggio et al., 1992; Schoups et al., 1995). A recent report utilizing *in vivo* calcium imaging of superficial cortical layers suggested that SRP requires locomotion for expression (Kaneko et al., 2017). However, abundant evidence demonstrates robust expression of SRP in layer 4 of the primary visual cortex (Aton et al., 2014; Cooke and Bear, 2010; Cooke et al., 2015; Frenkel et al., 2006; Gavornik and Bear, 2014; Kaplan et al., 2016).

The induction of experience-dependent plasticity in the cortex is also critically dependent on the state of the cortical network at the time of repetitive stimulation. Changes in the power of cortical oscillations have been frequently correlated with learning (Brickwedde et al., 2019; Howe et al., 2017; Park et al., 2016). Indeed, the power of high frequency oscillations (gamma) increases during short term memory (Jutras et al., 2009; Montgomery and Buzsáki, 2007) and working memory tasks (Lisman, 2010). Conversely, an increase in the ratio of theta to gamma power is associated with memory impairment (Moretti et al., 2009). In the visual cortex, the power of theta, alpha, beta and gamma oscillations increase in layer 4 following repetitive visual stimulation, leading to the intriguing suggestion that visual stimulus familiarity is encoded by changes in oscillatory power (Kissinger et al., 2018).

Finally, the response to repetitive visual stimulation is also modulated by inhibitory basket cells expressing the Ca^2+^ binding protein parvalbumin (PV INs). Suppression of PV IN excitability via optogenetic silencing or the NMDAR antagonist ketamine, inhibits SRP (Kaplan et al., 2016). Similarly, genetic deletion of the immediate early gene neuronal pentraxin 2 (NPTX2; aka NARP) which is highly enriched at excitatory synapses onto PV INs, inhibits SRP (Chang et al., 2010; O’Brien et al., 1999; Xu et al., 2003). Although NPTX2^−/−^ mice do not express SRP in response to LFVS, robust response potentiation can be induced with higher temporal frequency visual stimulation (HFVS, 10 Hz reversals /20 Hz screen refresh, Gu et al., 2013). This suggests that the different circuitry within the primary visual cortex is recruited by different temporal frequencies of repetitive visual stimulation.

Here we quantify the long-term impact of response potentiation induced in mouse primary visual cortex following low frequency and high frequency repetitive visual stimulation, defining the locus and specificity of the induced experience-dependent plasticity. We find that the robust SRP induced by LFVS that is typically limited to layer 4 of the visual cortex, extends to layer 2/3 following manipulations to enhance cortical plasticity. In contrast, HFVS induces robust and non-stimulus selective response potentiation in layers 4, and 5/6. The locus and the specificity of response potentiation is observed in response to visual stimuli that induce a phase reset of ongoing gamma oscillations.

## Results

### Control of the locus and specificity of visual response potentiation

Repetitive presentation of a simple visual stimulus at low temporal frequency over multiple days induces robust and stimulus-selective response potentiation (Aton et al., 2014; Cooke and Bear, 2010; Frenkel et al., 2006). To ask if an abbreviated visual stimulation protocol is sufficient to induce SRP, awake, head-fixed mice received truncated low frequency visual stimulation (LFVS; 200 presentations of full field, 100% contrast gratings, 0.05 cpd, 60 degree orientation reversing at 1 Hz /2 Hz screen refresh) and the visually-evoked response to familiar (60 degree) and novel (150 degree, 100% contrast gratings, 0.05 cpd) visual stimuli was quantified after 24 hours. Our induction protocol induced a significant increase in the VEP amplitude that was restricted to layer 4 of primary visual cortex and observed in response to the familiar, but not novel, visual stimulus (average VEP amplitude ± SEM in μV, initial: 100.52 ± 11.89; familiar: 123.08 ± 13.66; novel: 99.88 ± 12.07; n = 16 subjects. RANOVA _(df, 2, 15)_, F = 9.13, p < 0.001; initial v. familiar: p = 0.023; Bonferroni *post-hoc*, Fig. 1A). Simultaneously acquired single units, sorted into regular spiking (RS, presumptive excitatory) and fast spiking (FS, presumptive inhibitory), revealed a significant increase in the firing rate of RS neurons, also restricted to layer 4, in response to the familiar, but not novel, stimulus (average spike rate ± SEM in Hz, initial: 2.34 ± 0.21, n = 16 subjects, 33 units; familiar: 3.46 ± 0.44, n = 16, 28; novel: 2.83 ± 0.30, n = 16 subjects, 25 units. One way ANOVA _(df, 2, 83)_ F = 3.16, p = 0.047; initial v. familiar: p = 0.037; initial v. novel: p = 0.53; Tukey *post-hoc*).

**Figure 1.**
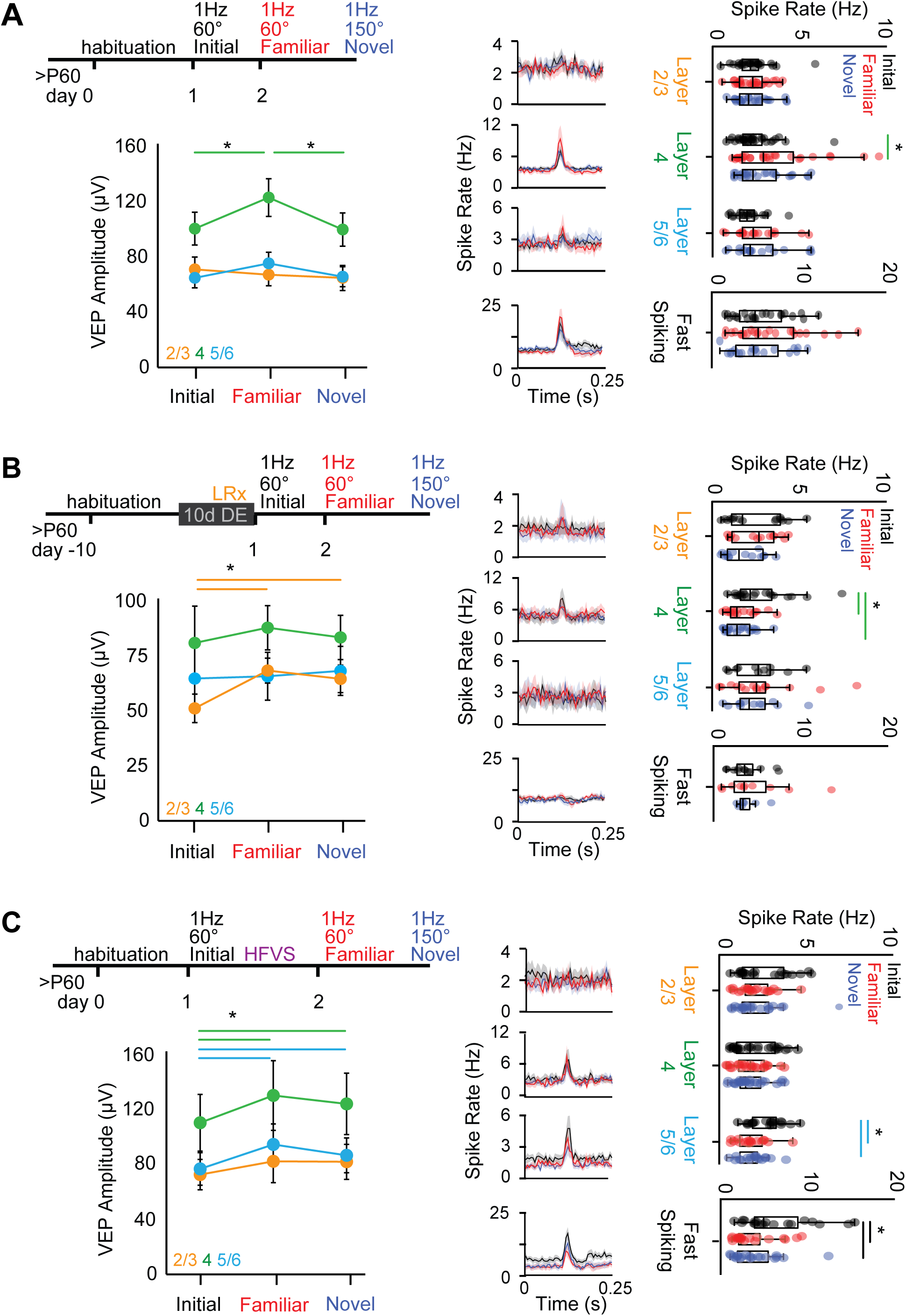
Visual experience regulates the location and generalizability of visual response potentiation. A) Left, inset: experimental timeline: subjects viewed 200 presentations of 0.05 cycle per degree, 100% contrast gratings, 60 degree orientation, reversing at 1 Hz (2 Hz screen refresh; initial LFVS). 24 hours later LFVS was repeated with the familiar orientation and a novel orientation. A significant increase in the amplitude of layer 4 VEPs in response to familiar, but not novel, visual stimulus was observed 24 hours after LFVS (RANOVA _(df, 2, 15)_, F= 9.13, p < 0.001; * = p < 0.05, Bonferroni *post hoc*; n = 16 subjects). Middle: peri-stimulus time histogram of average spike rate (0.05 ms bins), units sorted by layer (based on LFP waveform) and waveform shape (RS and FS). Right: Significant increase in average spike rates of RS layer 4 neurons in response to the familiar, but not novel, visual stimulus (ANOVA _(df, 2, 83)_, F = 3.16, p =0.047; * = p < 0.05, Tukey *post hoc*; RS n = 33 (initial) 28 (familiar) 25 (novel) units, 16 subjects). No change in layer 2/3 RS, layer 5/6 RS or FS firing rates. B) Left, inset: experimental timeline, as in A, except subjects received 10 days of DE and LRx prior to initial LFVS. Left, significant response potentiation of layer 2/3 VEP amplitudes in LRx subjects 24 hours after LFVS, in response to both familiar and novel visual stimuli (RANOVA _(df,2,10)_, F = 8.80, p = 0.001; * = p < 0.05, Bonferroni *post-hoc*; n = 11 subjects). Significant decrease in RS firing rates in layer 4 to familiar and novel stimuli (ANOVA _(df, 2, 55)_, F= 6.29, p = 0.003; * = p< 0.05, Tukey *post-hoc*; RS n = 20 (initial) 17 (familiar), 21 (novel) units, 11 subjects). No change in firing rates of layer 2/3 or 5/6 RS neurons or FS neurons. C) Left, inset: experimental timeline, as in A, except that high frequency visual stimulation (HFVS, 200 presentations of 0.05 cycle per degree 100% contrast gratings 30 degree orientation reversing at 10 Hz /20 Hz) was delivered following initial LFVS. Significant VEP potentiation in layers 4 (RANOVA _(df, 2, 10)_, F = 7.31, p = 0.004; * = p < 0.05, Bonferroni *post hoc*; n = 11 subjects) and 5/6 (RANOVA _(df, 2, 10)_, F = 11.41, p < 0.001; * = p < 0.05, Bonferroni *post hoc*; n = 11 subjects) in response to familiar and novel visual stimuli. HFVS induces a decrease in layer 5/6 RS spike rates to both familiar and novel stimuli (ANOVA _(df, 2, 54)_, F = 6.24, p = 0.003; * = p < 0.05, Tukey *post hoc*; RS n = 20 (initial), 17 (familiar), 20 (novel) units, 11 subjects), and a decrease in FS spiking rates (ANOVA _(df, 2, 56)_, F = 4.50, p = 0.015; * = p < 0.05, Tukey *post hoc*; n = 19 (initial), 20 (familiar), 20 (novel) units, 11 subjects).

The expression of several forms of synaptic plasticity that are typically limited to juveniles can be reactivated by light reintroduction (LRx) following dark exposure in adulthood (Gu et al., 2016; He et al., 2006; Huang et al., 2010; Montey and Quinlan, 2011). To ask if LRx regulates SRP, subjects received 10 days of dark exposure and LRx followed by the truncated LFVS protocol. LFVS induced a significant increase in the VEP amplitude that was restricted to layer 2/3, and observed in response to both familiar and novel visual stimuli (average VEP amplitude ± SEM in μV, initial: 44.66 ± 5.56; familiar: 59.44 ± 4.85; novel: 56.20 ± 5.36, n = 11 subjects. RANOVA _(df, 2, 10)_, F = 8.80, p = 0.001; initial v. familiar: p = 0.0142; initial v. novel: p = 0.044; Bonferroni *post-hoc*). Single unit recordings revealed a significant decrease in the firing rate of RS neurons, also restricted to layer 4, in response to both familiar and novel stimuli (average spike rate ± SEM in Hz, initial: 2.52 ± 0.32, n = 11 subjects, 20 units; familiar: 1.59 ± 0.19, n = 11 subjects, 17 units; novel: 1.433 ± 0.15, n = 11 subjects, 21 units. One way ANOVA _(df, 2, 55)_, F = 6.29, p = 0.003; initial v. familiar: p = 0.002; initial v. novel: p = 0.004; Tukey *post hoc*, Fig. 1B).

Increasing the temporal frequency of the repetitive visual stimulation expanded the locus and the selectivity of response potentiation. High frequency visual stimulation (HFVS, 200 presentations of full field, 100% contrast gratings, 0.05 cpd, 30 degree orientation, reversing at 10 Hz /20 Hz), induced a significant increase in VEP amplitudes in layers 4 and 5/6 in response to both familiar and novel visual stimuli observed after 24 hours (average VEP amplitude ± SEM in μV, layer 4: initial: 110.02 ± 19.34; familiar: 127.11 ± 22.82; novel: 125.33 ± 20.67, n = 11 subjects. RANOVA _(df, 2, 10)_, F = 7.31, p = 0.004; initial v. familiar: p = 0.038; initial v. novel: p = 0.005; Bonferroni *post-hoc*. Layer 5/6: initial: 80.80 ± 10; familiar: 97.93 ± 13.75; novel: 91.43 ± 11.73, n = 11 subjects. RANOVA _(df, 2, 10)_, F = 11.41, p < 0.001; initial v. familiar: p = 0.005; initial v. novel: p = 0.022; Bonferroni *post-hoc*. Fig. 1C). A generalized VEP potentiation was induced 24 hour later if HFVS was presented after (Fig 1C) or prior to LFVS (Fig. S1). Potentiated VEP amplitude was also seen in response to visual stimuli of novel spatial frequencies (Fig. S2). Interestingly, HFVS induced a significant decrease in the visually evoked firing rates of FS interneurons (average spike rate ± SEM in Hz, initial: 7.00 ± 0.98, n = 11 subjects, 19 units; familiar: 4.071 ± 0.59, n = 11 subjects, 20 units; novel: 4.39 ± 0.64, n =11 subjects, 20 units. ANOVA _(df, 2, 57)_ F = 4.50, p = 0.015; initial v. familiar: p = 0.022; initial v. novel: p = 0.046; Tukey *post hoc*) and layer 5/6 RS neurons (average spike rate ± SEM in Hz, initial: 2.35 ± 0.18, n = 11 subjects, 20 units; familiar: 1.62 ± 0.19, n = 11 subjects, 17 units; novel: 1.59 ± 0.15 Hz, n = 11 subjects, 20 units. One way ANOVA _(df, 2, 54)_, F = 6.24 p = 0.003; initial v. familiar: p = 0.014; initial v. novel: p = 0.007; Tukey *post-hoc*). In contrast, repetitive presentation of visual stimuli at other temporal frequencies (5 Hz/10 Hz or 20 Hz/40 Hz) did not induce response potentiation (Fig. S3). Indeed, HFVS induced more widespread c-fos expression and higher visually-evoked firing in RS neurons than other temporal frequencies (Fig. S4).

### Visual stimulation modifies ongoing cortical oscillation power and phase

Cortical oscillatory power and phase can be entrained by incoming sensory stimulation and alter the response to subsequent visual stimulation (Cardin et al., 2009; Kissinger et al., 2018). To ask how repetitive visual stimulation impacts ongoing cortical oscillatory activity, we examined the average absolute value of the time-locked analytic signal during visual stimulation relative to pre-stimulation spontaneous activity (evoked by equal illuminant grey screen; 26 cd/m^2^). No coincident changes in oscillatory power in any cortical layer were observed during LFVS (Fig. 2A), while LFVS following LRx induced an increase in beta power (13 - 30 Hz) within layer 2/3 (% change in power ± SEM, one-sided t-test, 4.57 ± 1.76, n = 11 subjects, t= 2.19, p = 0.03. Fig. 2C). In contrast, HFVS induced a significant decrease in gamma power (30 - 100 Hz) in all layers (% change in power ± SEM, one sided t-test. Layer 2/3: −9.52 ± 2.41, n = 11 subjects, t = 3.52, p = 0.002. Layer 4: −8.45 ± 2.09, n = 11 subjects, t = 3.62, p = 0.002. Layer 5/6: −6.02 ± 2.23, n = 11 subjects, t = 2.29, p = 0.023. Fig. 2E). Widespread changes in cortical power in all layers of primary visual cortex were also observed during lower and higher frequency stimulation (Fig. S5A, B).

**Figure 2.**
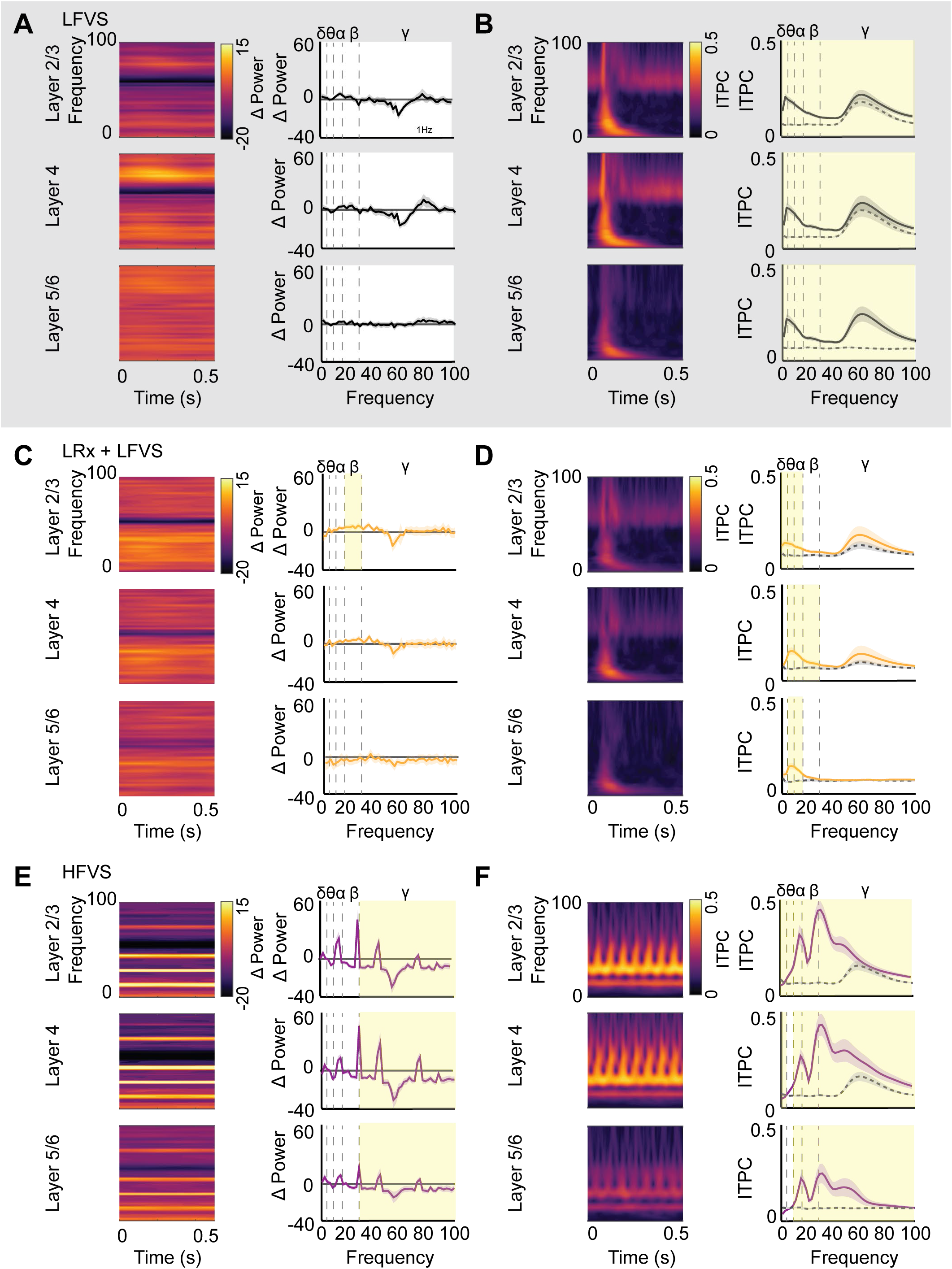
Oscillatory power and visually evoked phase reset during LFVS and HFVS. Left; heat map depicts average oscillatory power from 0 to 100 Hz (3 Hz bins) across cortical layers for first 0.5 seconds of each trial, calculated from the analytic signal of the LFP by layer. Power is normalized spontaneous cortical activity in response to 26 cd/m^2^ grey screen. Average power over entire trial is binned within each frequency band (delta: 1 – 4, theta: 4 - 8, alpha: 7 - 13, beta: 13 - 30, gamma: 30 – 100 Hz). Left; ITPC from 0 to 100 Hz (in 3 Hz bins) for 0.5 seconds of each trial, calculated using Morlet wavelet convolution. Average ITPC over entire trial. A) No change in average oscillatory power in any cortical layer during initial LFVS in naïve subjects (n = 16 subjects). B) ITPC reveals significant increase in visual phase reset in high and low frequency bands across all cortical layers during LFVS (paired t-test, yellow highlight = p < 0.05; n = 16 subjects). C) LFVS following DE and LRx increased beta power in layer 2/3 only (one-sided t-test, yellow highlight = p < 0.05; n = 11 subjects). D) Significant increase in phase reset of low frequency oscillations in all cortical layers during LFVS after LRx (paired t-test, yellow highlight = p < 0.05; n = 11 subjects). E) HFVS evokes a significant decrease in average gamma power in all cortical layers (one sided t test, yellow highlight = p < 0.05; n = 11 subjects). F) HFVS induces significant changes in ITPC throughout the cortical depth, including a significant increase in visually driven phase reset in all frequency bands alpha and above for all layers and a significant decrease in the delta (paired t-test, yellow highlight = p < 0.05; n = 11 subjects).

To ask if repetitive visual stimulation impacted the phase of ongoing oscillatory activity, we convolved the LFP signal with a complex Morlet wave and calculated the time-locked angle of the resultant complex output. Inter-trial phase consistency (ITPC) was calculated from the time-locked phase, ranging from 0 (if oscillatory phase is random and not reset by incoming visual input) and 1 (if oscillatory phase was similarly reset in all trials). LFVS reset the phase of all oscillatory bands throughout the cortical depth during visual stimulation (Fig. 2B, Table 1). Following LRx, LFVS continued to reset low, but not high, frequency oscillations (Fig. 2D). HFVS significantly increased ITPC in all cortical layers to all frequencies above alpha (7 -13 Hz, Fig. 2F).

**Table 1.** Mean and standard deviation for values in Figure 2 B, D, & F.

|  | Layer |  | Delta<br>(1- 4 Hz) | Theta<br>(4-8 Hz) | Alpha<br>(7-13 Hz) | Beta<br>(13-30 Hz) | Gamma<br>(30-100 Hz) |
| --- | --- | --- | --- | --- | --- | --- | --- |
| LFVS | 2/3<br>n=15 | Spontaneous | 0.068±0.003 | 0.066±0.004 | 0.065±0.003 | 0.065±0.002 | 0.111±0.010 |
|  |  | Stimulation | 0.163±0.013 | 0.183±0.015 | 0.153±0.011 | 0.113±0.006 | 0.137±0.010 |
|  |  | p value | <0.001* | <0.001* | <0.001* | <0.001* | 0.035* |
|  | 4<br>n=16 | Spontaneous | 0.063±0.003 | 0.062±0.003 | 0.063±0.002 | 0.064±0.002 | 0.126±0.018 |
|  |  | Stimulation | 0.168±0.013 | 0.199±0.020 | 0.165±0.012 | 0.117±0.009 | 0.157±0.017 |
|  |  | p value | <0.001* | <0.001* | <0.001* | <0.001* | 0.034* |
|  | 5/6<br>n=15 | Spontaneous | 0.070±0.006 | 0.063±0.002 | 0.063±0.002 | 0.063±0.001 | 0.062±0.0015 |
|  |  | Stimulation | 0.122±0.011 | 0.166±0.016 | 0.142±0.011 | 0.099±0.007 | 0.084±0.005 |
|  |  | p value | 0.001* | <0.001* | <0.001* | 0.001* | <0.001* |
| LRx +LFVS | 2/3<br>n=11 | Spontaneous | 0.071±0.008 | 0.069±0.004 | 0.066±0.005 | 0.067±0.005 | 0.089±0.010 |
|  |  | Stimulation | 0.127±0.015 | 0.136±0.016 | 0.117±0.011 | 0.093±0.008 | 0.118±0.021 |
|  |  | p value | 0.018* | 0.006* | 0.001* | 0.061 | 0.223 |
|  | 4<br>n=11 | Spontaneous | 0.070±0.008 | 0.065±0.004 | 0.066±0.003 | 0.070±0.005 | 0.079±0.006 |
|  |  | Stimulation | 0.098±0.009 | 0.157±0.019 | 0.143±0.013 | 0.097±0.011 | 0.103±0.019 |
|  |  | p value | 0.08 | <0.001* | <0.001* | 0.070 | 0.270 |
|  | 5/6<br>n=10 | Spontaneous | 0.085±0.015 | 0.061±0.004 | 0.063±0.003 | 0.071±0.005 | 0.070±0.005 |
|  |  | Stimulation | 0.100±0.015 | 0.162±0.017 | 0.129±0.13 | 0.084±0.008 | 0.070±0.004 |
|  |  | p value | 0.547 | 0.001* | 0.001* | 0.287 | 0.932 |
| HFVS | 2/3<br>n=11 | Spontaneous | 0.079±0.007 | 0.065±0.003 | 0.066±0.003 | 0.069±0.002 | 0.103±0.011 |
|  |  | Stimulation | 0.051±0.003 | 0.094±0.010 | 0.186±0.028 | 0.292±0.039 | 0.180±0.025 |
|  |  | p value | 0.011* | 0.007* | <0.001* | <0.001* | 0.014* |
|  | 4<br>n=11 | Spontaneous | 0.076±0.007 | 0.066±0.0045 | 0.067±0.004 | 0.066±0.003 | 0.099±0.011 |
|  |  | Stimulation | 0.051±0.004 | 0.086±0.010 | 0.150±0.018 | 0.253±0.030 | 0.193±0.039 |
|  |  | p value | 0.035* | 0.062 | <0.001* | <0.001* | 0.025* |
|  | 5/6<br>n=11 | Spontaneous | 0.070±0.011 | 0.067±0.006 | 0.066±0.003 | 0.062±0.003 | 0.063±0.001 |
|  |  | Stimulation | 0.050±0.003 | 0.071±0.006 | 0.124±0.011 | 0.172±0.019 | 0.113±0.021 |
|  |  | p value | 0.152 | 0.572 | <0.001* | <0.001* | 0.036 |

### Changes in the power of spontaneous cortical oscillations emerges following visual stimulation

To ask if truncated visual stimulation induced in a change in the distribution of spontaneous cortical oscillatory activity, we quantified spontaneous oscillations and single unit activity 24 hours after visual stimulation. Following LFVS a significant increase in alpha power (7 - 13 Hz) in all cortical layers and increased beta (13 - 30 Hz) power in layer 5/6 emerged (% change in alpha ± SEM, one-sided t-test: layer 2/3: 2.21 ± 0.78, n = 15 subjects. t = 2.44, p = 0.014. Layer 4: 2.31 ± 0.89, n = 16 subjects; t = 2.42, p = 0.020. Layer 5/6: 3.50 ± 0.95, n = 15 subjects; t = 3.28, p = 0.002; % change in beta: layer 5/6: 3.24 ± 1.21, n = 15 subjects; t = 2.29, p = 0.018; Fig. 3A). However, we observed no changes in spontaneous firing rates of either RS or FS neurons 24 hours after LFVS. Following LFVS to LRx subjects, the power of spontaneous theta oscillations decreased in layer 5/6 (% change in power ± SEM, one sided t-test: −6.82 ± 2.16, n = 10 subjects, t = 2.03, p = 0.034), but there was no change in SU firing rates (Fig. 3B). In contrast, 24 hours after HFVS we observed a significant increase in spontaneous oscillatory power in all cortical layers at all frequency bands above alpha (7 - 13 Hz) and a decrease in delta power (1 - 4 Hz; Table 2). A significant decrease in spontaneous firing rates of FS interneurons (average spike rate ± SEM in Hz, day 1: 5.91 ± 0.88, n = 11 subjects, 19 units; day 2: 3.12 ± 0.51, n = 11 subjects, 18 units. Student’s t-test, t= 2.41, p = 0.011) and layer 5/6 RS neurons was observed 24 hours after visual stimulation (average spike rate ± SEM in Hz, spontaneous day 1: 2.39 ± 0.15, n = 11 subjects, 20 units; day 2: 1.56 ± 0.22, n = 11 subjects, 15 units. Student’s t-test, t = 2.63, p = 0.006. Fig. 3C). Thus wide-spread changes in oscillatory power and spontaneous neuronal activity emerge following HFVS.

**Table 2.** Mean and standard deviation for values in Figure 3 C.

|  | Layer |  | Delta<br>(1- 4 Hz) | Theta<br>(4-8 Hz) | Alpha<br>(7-13 Hz) | Beta<br>(13-30 Hz) | Gamma<br>(30-100 Hz) |
| --- | --- | --- | --- | --- | --- | --- | --- |
| HFVS<br>n = 11 | 2/3 | Percent Change | -9.47±2.25 | 2.22±1.21 | 1.77±0.76 | 4.07±1.57 | 6.40±2.21 |
|  |  | p value | 0.001* | 0.096 | 0.043* | 0.027* | 0.015* |
|  | 4 | Percent Change | -9.99±2.15 | 2.28±1.08 | 2.43±1.052 | 4.82±1.78 | 6.47±2.51 |
|  |  | p value | <0.001* | 0.016* | 0.072 | 0.016* | 0.036* |
|  | 5/6 | Percent Change | -9.62±2.11 | 1.89±1.125 | 3.08±1.27 | 5.29±1.31 | 6.11±1.98 |
|  |  | p value | 0.001* | 0.162 | 0.035* | 0.002* | 0.011* |

**Figure 3.**
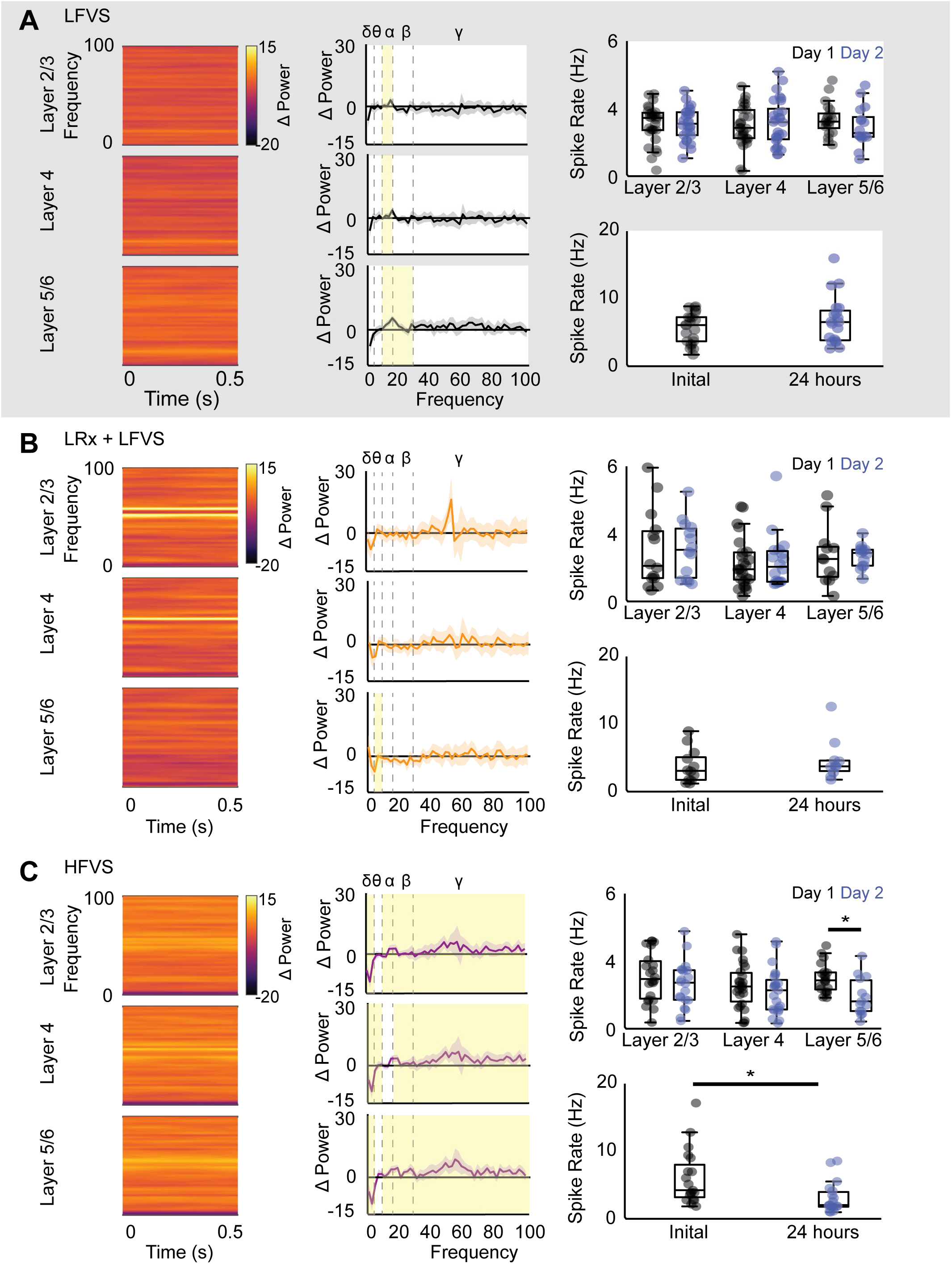
Sustained changes in oscillatory power and spontaneous firing rates induced by HFVS. Left: heat map, average change in spontaneous oscillatory power 24 hours after visual stimulation (percent change from pre-experimental spontaneous). Middle: average power during 1 second epoch. Right: top: spontaneous RS firing rates by layer and time point, bottom: spontaneous FS firing rates by time point. A) A significant increase in spontaneous alpha power is observed in all cortical layers and an increase in beta power in layer 5/6 24 hours after LFVS (one-sided t-test, yellow highlight = p < 0.05; n = 15 subjects). No change in spontaneous firing rates of RS or FS units. B) A significant increase in spontaneous theta power 24 hours after LFVS following LRx in layer 5/6 (one sided t-test, yellow highlight = p < 0.05; n = 10 subjects) with no change in spontaneous firing rates of RS or FS units. C) A significant increase in high and mid frequency oscillatory power in all cortical layers, as well as a significant decrease in delta power in all layers 24 hours after HFVS (one sided t-test, yellow highlight = p < 0.05; n = 11 subjects). A significant decrease in spontaneous firing rates of layer 5/6 RS neurons (unpaired t-test, * = p < 0.05; n = 20 (day 1), 15 (day 2) units, 11 subjects and FS neurons 24 hours after HFVS (unpaired t-test, * = p < 0.05; n = 19 (day 1), 18 (day 2) units, 11 subjects.

### Change in oscillatory power does not predict expression of response potentiation

To ask if visually-evoked changes in oscillatory power predict response potentiation 24 hours later, we examined visually evoked power in response to the familiar and novel visual stimulus in the temporal window of maximal visually evoked activity after stimulus onset (0 - 100 ms). LFVS induced a significant increase in alpha and beta power in response to the familiar visual stimulus in layer 4, coincident with the expression of SRP. However, an increase in alpha and beta power was also observed in response to familiar and novel visual stimuli in layer 5/6 (% change in power ± SEM, RANOVA _(df, 2, 15)_ with Bonferroni *post hoc*. Layer 4 alpha: initial: 0.63 ± 1.50; familiar: 4.51 ± 1.32, n = 16 subjects, F = 8.36, p = 0.001; initial v. familiar, p = 0.001; beta: initial: 2.20 ± 2.38; familiar: 6.67 ± 3.03, n = 16 subjects, F=4.10, p = 0.026; initial v. familiar: p = 0.007. Layer 5/6 alpha: initial: 0.32 ± 1.36; familiar: 4.62 ± 1.18; novel: 4.44 ± 1.54, n = 15 subjects, F = 21.26, p < 0.001; initial v. familiar: p < 0.001; initial v. novel: p < 0.001; beta: initial: 0.43 ± 1.53; familiar: 5.26 ± 1.96; novel: 3.32 ± 1.39, n = 15 subjects, F=6.55, p = 0.004; initial v. familiar: p = 0.005; initial v. novel: p = 0.034, Fig. 4A). In LRx subjects, we observed no change in oscillatory power in any cortical layers following LFVS (Fig. 4B), despite robust generalized response potentiation in layer 2/3. In contrast, HFVS induced a significant increase in beta power in all cortical layers in response to familiar and novel visual stimuli (% change in power ± SEM, RANOVA _(df, 2, 10)_ with Bonferroni *post-hoc*. Layer 2/3: initial: 0.22 ± 1.03; familiar: 7.28 ± 2.23; novel: 5.80 ± 1.65, n = 11 subjects, F = 9.82, p = 0.001; initial v. familiar: p = 0.016; initial v. novel: p = 0.006. Layer 4: initial: 2.29 ± 1.95; familiar: 10.96 ± 3.64; novel: 8.04 ± 2.33, n = 11 subjects, F = 10.25, p < 0.001; initial v. familiar: p = 0.014; initial v. novel: p = 0.003. Layer 5/6: initial: −0.33 ± 0.66; familiar: 7.18 ± 1.77; novel: 5.22 ± 1.15, n = 11 subjects, F = 10.45, p < 0.001; initial v. familiar: p = 0.007; initial v. novel: p = 0.016, Fig. 4C). Together these results indicate that the changes in oscillatory power induced by visual experience do not predict the spatial distribution or stimulus specificity of the observed response potentiation.

**Figure 4.**
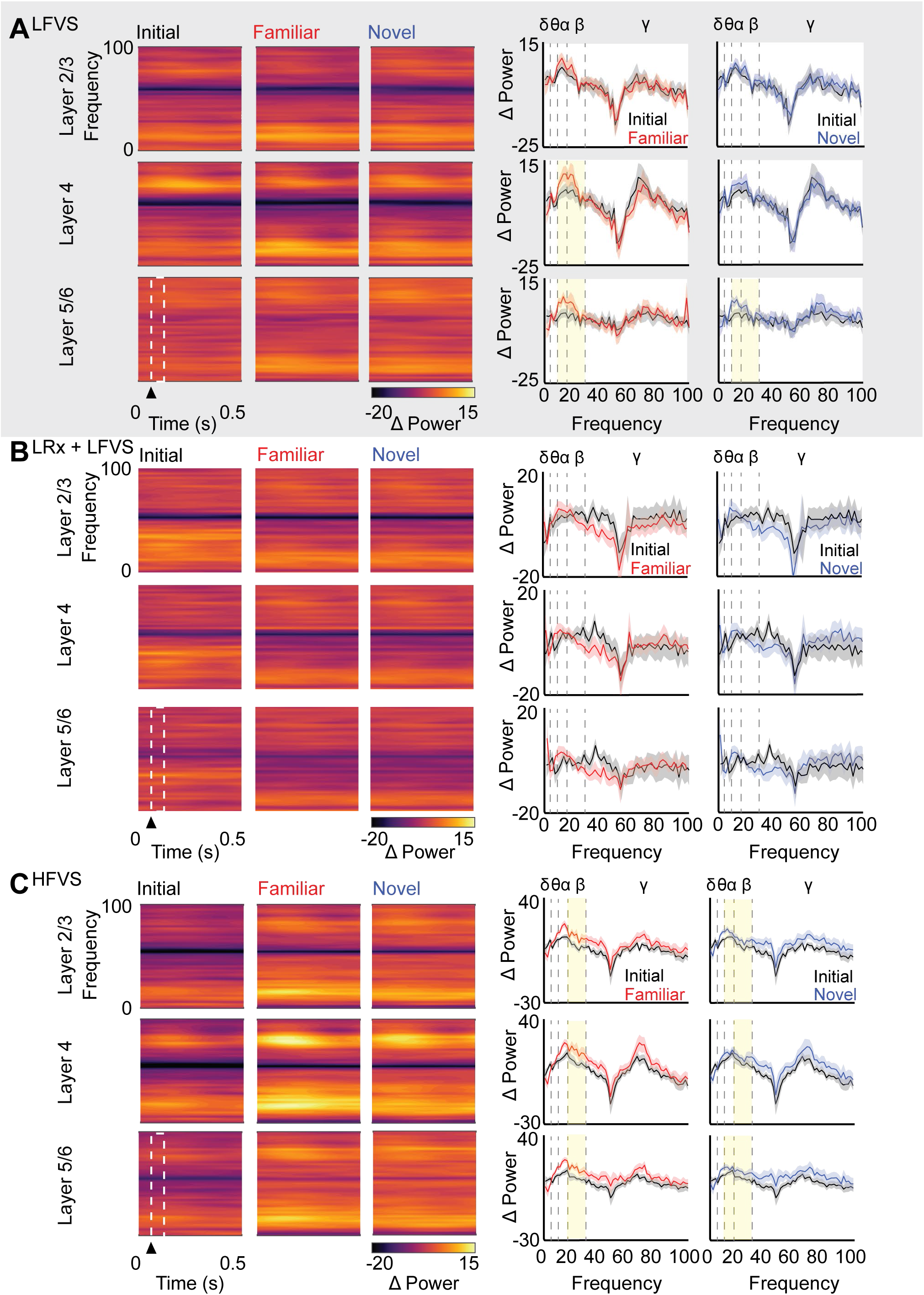
Visually-evoked changes in oscillatory power do not predict VEP potentiation. Left: heat map: normalized trial average oscillatory power during visual stimulation, arrowhead indicates onset of visual stimulus. Right: peak cortical oscillatory power during 100 ms after stimulus onset. White box on heat map denotes time slice. A) In layer 4, a significant increase in alpha and beta power in response to familiar visual stimulus 24 hours after LFVS (RANOVA _(df, 2, 15)_, with Bonferroni *post hoc*; n = 16 subjects: alpha: F = 8.36, p = 0.001, beta: F = 4.10, p = 0.026; yellow highlight = p < 0.05). In layer 5/6, a significant increase in alpha and beta power in response to both familiar and novel visual stimulus (RANOVA _(df, 2, 14)_, with Bonferroni *post hoc*; n = 15 subjects: alpha: F = 21. 26, p < 0.001, beta: F = 6.55, p = 0.004; yellow highlight = p < 0.05). B) No change in instantaneous power in response to familiar or novel visual stimulus in any layer 24 hours after LFVS in LRx subjects (n = 11 subjects). C) A significant increase in beta power in response to novel and familiar visual stimulus 24 hours after HFVS (RANOVA _(df, 2, 10)_, with Bonferroni *post-hoc*; n = 11 subjects: layer 2/3: F = 9.82, p = 0.001, layer 4: F = 10.25, p < 0.001, layer 5: F = 10.45, p < 0.001; yellow highlight = p < 0.05).

### Oscillatory phase predicts the expression of response potentiation

Changes in oscillatory power in the absence of phase synchronization would increase response variability, rather than response magnitude. We therefore hypothesized that the increase in response magnitude induced by repetitive visual stimulation is mediated by a reset of oscillatory phase (Voloh and Womelsdorf, 2016). To test this hypothesis, we calculated the average ITPC in the temporal window that corresponds to maximal visually evoked activity after stimulus onset (0 - 100 ms), 24 hours after repetitive visual stimulation. LFVS induced a significant increase in beta and gamma ITPC in cortical layer 4 that was restricted to the familiar visual stimulus, coincident with the expression of SRP (Fig 5A, average ITPC ± SEM, RANOVA _(df, 2, 15)_ with Bonferroni *post-hoc*, beta: initial: 0.36 ± 0.047; familiar: 0.47 ± 0.052, n = 16 subjects, F = 5.20, p = 0.011; initial v. familiar: p = 0.045. Gamma: initial: 0.26 ± 0.029; familiar: 0.35 ± 0.043, n = 16 subjects, F = 8.82, p < 0.001; initial v. familiar: p = 0.005). Similarly, following LRx, LFVS significantly increased gamma ITPC in layer 2/3 in response to familiar and novel visual stimuli, mirroring the expression of response potentiation (Fig 5B, initial: 0.15 ± 0.023; familiar: 0.28 ± 0.027; novel: 0.25 ± 0.023, n = 11 subjects. RANOVA _(df, 2, 10)_, F = 11.41, p < 0.001; initial v. familiar: p = 0.006, initial v. novel: p = 0.005; Bonferroni *post-hoc*). Finally, HFVS significantly increased gamma ITPC in all cortical layers, in response to both familiar and novel visual stimuli, coincident with the observed widespread response potentiation (Fig 5C, RANOVA _(df, 2, 10)_ with Bonferroni *post-hoc*, layer 2/3: initial: 0.18 ± 0.027; familiar: 0.32 ± 0.039; novel: 0.29 ± 0.038, n = 11 subjects, F = 19.42, p < 0.001; initial v. familiar: p = 0.001; initial v. novel: p = 0.003. Layer 4: initial: 0.20± 0.036; familiar: 0.30 ± 0.042; novel: 0.30 ± 0.045, n = 11 subjects, F = 13.04, p < 0.001; initial v. familiar: p = 0.010; initial v. novel: p = 0.006. Layer 5: initial: 0.16 ± 0.036; familiar: 0.24 ± 0.038; novel: 0.21 ± 0.040, n = 11 subjects, F = 5.29, p = 0.025; initial v. familiar: p = 0.022; initial v. novel: p = 0.021). Together this suggests that the strength of oscillatory phase reset in response to incoming visual stimulus predicts the magnitude, location and specificity of response potentiation.

**Figure 5.**
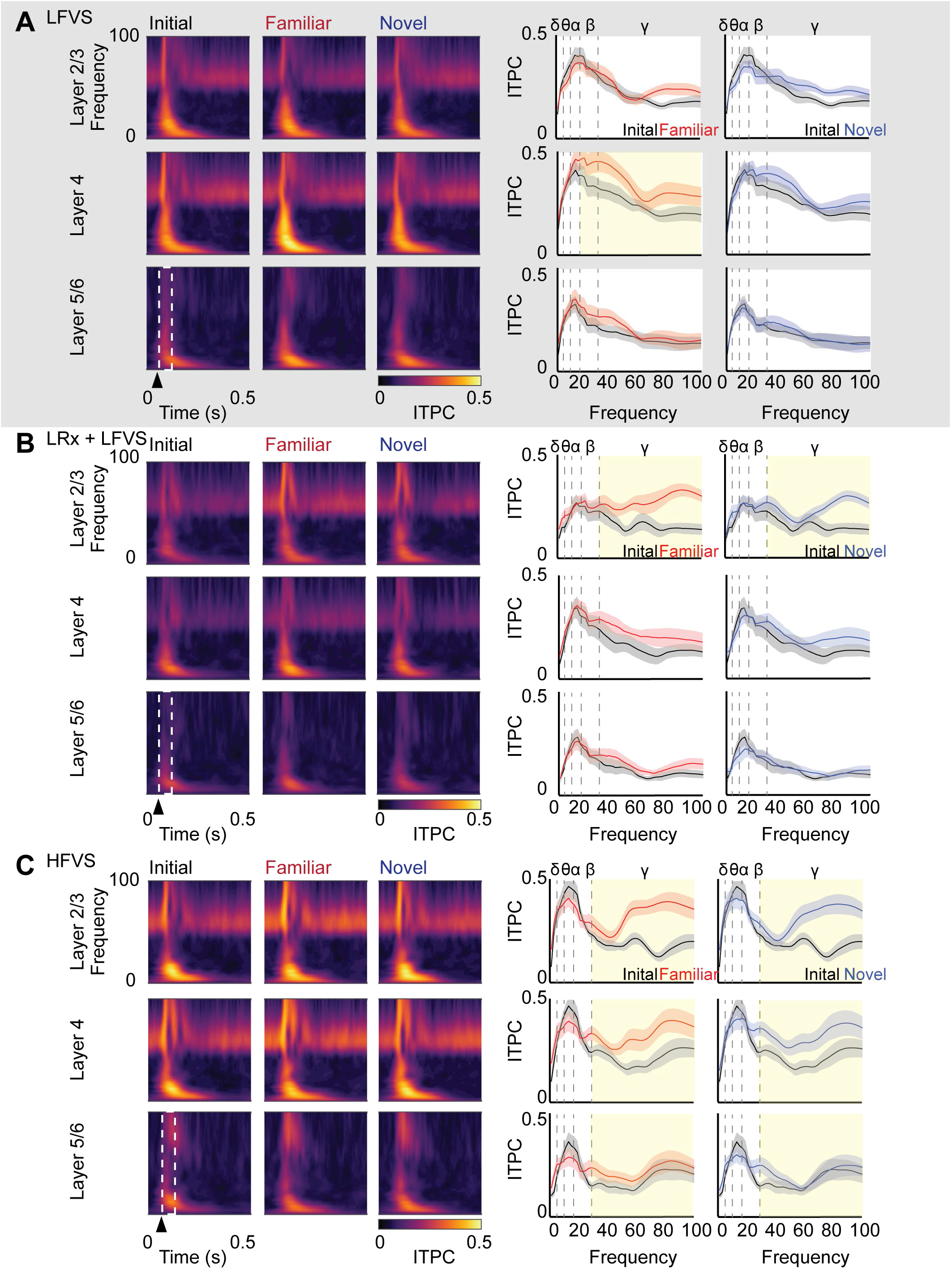
Visually-evoked phase reset predicts VEP potentiation. Left: heat map: peak ITPC from 0 - 100 Hz. Right: average of the first 100 ms after stimulus onset. White box in heat map (left) denotes time slice. A) Significant increase in layer 4 oscillatory phase reset of beta and gamma oscillations in response to familiar stimulus 24 hours after LFVS (RANOVA _(df, 2, 15)_, with Bonferroni *post-hoc*; n =16 subjects: beta: F = 5.20, p = 0.011, gamma: F = 8.82, p < 0.001; yellow highlight = p < 0.05). B) A significant increase in the evoked reset of gamma oscillatory phase in layer 2/3, in response to both familiar, and novel visual stimuli 24 hours after LFVS in LRx subjects (RANOVA _(df,2,10)_, Bonferroni *post-hoc*; n = 11 subjects, F = 4.41, p < 0.001; yellow highlight = p < 0.05). C) A significant increase in the evoked reset of gamma oscillatory phases in layers 2/3, 4, and 5/6 in response to familiar and novel visual stimuli, 24 hours after HFVS (RANOVA _(df, 2, 10)_, Bonferroni post-hoc, n = 11 subjects: layer 2/3: F = 19.42, p < 0.001, layer 4: F = 13.04, p < 0.001, layer 5: F = 5.29, p = 0.025; yellow highlight = p < 0.05).

## Discussion

Stimulus-selective response potentiation and visual perceptual learning can improve sensory detection thresholds and sensory discrimination in the aged or damaged cortex (Camilleri et al., 2014; Eaton et al., 2016; Levi and Li, 2009; Sale and Berardi, 2015). Repetitive presentation of a visual stimulus or repetitive performance in a visual discrimination task for hundreds of trials over many days is typically used to induce SRP and perceptual learning. Here we show that a truncated protocol of only 200 repetitions of a simple visual stimulus is sufficient to induce sustained changes in primary visual cortex. The rapid SRP induced by LFVS is expressed as a potentiation of VEP amplitude and an increase in RS neuron activity that is restricted to layer 4 of primary visual cortex and is selective for the familiar stimulus. In contrast, LFVS delivered to adults following LRx after dark exposure, enabled response potentiation in layer 2/3 to both familiar and novel stimuli. The loss of SRP in layer 4 is interesting, but not unexpected, as LRx induces an increase in the activity of the extracellular protease MMP9 at thalamo-cortical synapses and is proposed to decrease feedforward excitation to cortical layer 4 (Murase et al., 2017). SRP, although typically absent in superficial layers, is also observed in supragranular cortical layers following the enhancement of plasticity in primary visual cortex by locomotion (Kaneko et al., 2017). The locus of expression as well as the generalizability of response potentiation is regulated by the temporal frequency of repetitive visual stimulation, as HFVS induces a non-selective potentiation of VEP amplitudes in layers 4 and 5/6. A sustained decrease in the visually-evoked and spontaneous activity of FS interneurons in primary visual cortex was also induced by HFVS, and a counter-intuitive decrease in the firing rates of layer 5/6 RS, which may further reduce inhibition via long-range translaminar axons (Bortone et al., 2014; Olsen et al., 2012).

SRP of VEP amplitudes and spiking output of RS neurons emerge slowly over days (Frenkel et al., 2006) and require sleep for consolidation (Aton et al., 2014), mimicking some of the key features of other forms of perceptual learning (Alain et al., 2015; Karni et al., 1994). Indeed, we observe no difference in VEP amplitude immediately following HFVS. The slow emergence of response potentiation contrasts with many other forms of experience-dependent synaptic plasticity, such as the response to photic tetanus (Clapp et al., 2006; Teyler et al., 2005) and acute response adaptation to repetitive stimulation (Taaseh et al., 2011). Interestingly, acute potentiation of VEP amplitude can be induced during the acquisition of the steady state responses by directed attention or interjection of a deviant visual stimulus, i.e. grating rotated from familiar (Joon Kim et al., 2007; Morgan et al., 1996). This potentiation is attributed to an increase in visual response reset coincident with the visual stimulus, suggesting visually-evoked changes in cortical oscillatory synchrony.

Learning is also frequently correlated with changes in oscillatory power (Brickwedde et al., 2019; Howe et al., 2017; Park et al., 2016) Recent work suggests an increase in the power of low frequency oscillations emerge in rodent cortex during repetitive visual stimulation, which may encode the familiarity of the stimulus (Kissinger et al., 2018). An increase in beta frequency power and beta/gamma coupling in visual thalamus and primary visual cortex (beta: 16 - 24, gamma: 30 - 45) is also observed in cats after Pavlovian conditioning to associate a reward with a visual, but not auditory, stimulus (Bekisz and Wróbel, 1999). Following HFVS, all visual stimuli induce an increase in beta power (13 - 30 Hz), suggesting that the visual cortex may perceive all stimuli as familiar. However, changes in oscillatory power in the absence of phase synchronization would result in an increase in response variability, not magnitude, and are insufficient to predict the locus or specificity of response potentiation. Here we used ITPC to track interactions between the time-locked visual stimulation and ongoing oscillatory activity in primary visual cortex. We found that visually-evoked oscillatory phase reset and the synchrony of gamma are highly predictive of the locus and specificity of visual response potentiation: 1) Following LFVS, visually-evoked phase reset of gamma oscillations and SRP were restricted to the familiar stimulus and layer 4. 2) LFVS following LRx induced phase reset and SRP in response to familiar and novel stimuli in layers 2/3. 3) HFVS induced a visually-evoked phase reset of gamma oscillations in all layers and in response to novel and familiar visual stimuli, coincident with the majority of response potentiation. Oscillatory phase reset may be a fundamental mechanism of potentiation of sensory responses. Indeed, synchronization of the phase of a fluctuating visual stimulus with ongoing oscillatory activity is known to increase the LFP amplitude (Kissinger et al., 2018), and the well-documented increase sensory response amplitudes following directed attention or in response to a deviant stimulus are coincident with an oscillatory phase reset (Fuentemilla et al., 2008; Heinze et al., 1994; Hillyard and Anllo-Vento, 1998; Stothart and Kazanina, 2013; Zhang and Luck, 2009).

Our data support a model in which HFVS promotes generalized response potentiation by inducing a sustained change in the oscillatory activity of primary visual cortex and an increase in the sensitivity to oscillatory phase reset by subsequent visual stimulation. Although the characteristics that regulate vulnerability to phase reset are unknown, one possibility is that the higher frequency gamma oscillations induced by HFVS are more sensitive to phase reset. Gamma oscillations are an interesting candidate for entraining visual responses as cortical gamma is thought to be generated within feedforward thalamo-cortical connections (Bastos et al., 2015; van Kerkoerle et al., 2014; Spaak et al., 2012), but the strength and phase synchrony of gamma are modulated/entrained by the output of cortical PV INs. The reduction in the excitability of PV INs following HFVS may reduce cortical PV IN modulation of gamma oscillations, thereby enabling entrainment to visual stimulation. Indeed, optogenetic silencing of PV IN output during visual stimulation increases oscillatory synchrony with single unit activity across multiple frequencies, including gamma (Chen et al., 2017). In contrast, optogenetic activation of PV INs in barrel cortex induces a change in the peak frequency of gamma oscillations and blunts the response to sensory stimulation, showing that gamma oscillation regulation of the response to sensory stimulation is modulated by PV INs (Cardin et al., 2009). Driving PV INs at gamma frequency using dynamic-clamp reduces response reliability of RS neurons, consistent with the prediction that change in oscillatory power in the absence of oscillatory phase reset increases response variability (Sohal et al., 2009).

HFVS stimulation appeared to be optimal for the induction of gamma oscillations, and for evoking higher firing rates in RS layer 4 neurons than visual stimulation at other temporal frequencies. This was surprising given that the preferred temporal frequency for drifting gratings observed in anesthetized and awake mice is 2 Hz (Niell and Stryker, 2008), however higher temporal frequencies are above flicker fusion in the murine visual system (Durand et al., 2016; Porciatti et al., 1999; Tanimoto et al., 2015). Our use of high frequency repetitive visual stimulation to induce brain wave entrainment and enhancement of sensory response strength was motivated by recent work in humans in which experiments in humans in which high frequency visual stimulation (20 Hz flicker of elongated bars) induced a long-lasting improvement in orientation discrimination (Beste et al., 2011; Marzoll et al., 2018). Exposure to a 10 Hz flickering visual stimulation presented during a word recognition task also improved performance (Williams, 2001; Williams et al., 2006). In contrast, low frequency stimulation (1 Hz visual stimulus) did not impact visual discrimination (Marzoll et al., 2018). Trans-cranial brain stimulation techniques including direct current stimulation (tDCS), alternating current stimulation (tACS), and repetitive transcranial magnetic stimulation (rTMS) also alter performance on sensory and memory tasks. However the effects of trans-cranial brain stimulation are relatively short-lived (seconds-minutes) and susceptible to disruption (Albouy et al., 2017; Hanslmayr et al., 2019; Hordacre et al., 2015; Huang et al., 2009; Todd et al., 2009). Interestingly, in mice, rTMS paired with an oriented visual stimulus increases the visually evoked calcium responses to familiar and novel stimuli. However, the generalized response potentiation required repeated rTMS+visual stimulus pairings over several days (Tsapa et al., 2019). In contrast, the effects of LFVS and HFVS in rodents are evident 24 hours after stimulation and can be induced by a single bout of visual stimulation. The generalizable response enhancement observed after truncated HFVS suggests a potential role in rapid, non-invasive vision therapy.

## Methods

### Animals

Equal numbers of adult (postnatal day 60 - 90) male and female C57BL/6J mice (Jackson Lab, Bar Harbor, ME) were utilized. Animals were housed on a 12:12 hour dark:light cycle with food and water *ad libitum*, and experiments were initiated ~ 6 hours into the light phase. Animals assigned to the light reintroduction (LRx) group were housed in a light tight room for 10 days, with daily care provided under infrared light. All procedures conformed to the guidelines of the University of Maryland Institutional Animal Care and Use Committee. Sample sizes were determined by previous studies quantifying the effect of visual experience on visual response amplitudes.

### Electrophysiology

House-made 1.2 mm length 16-channel shank electrodes were implanted into binocular primary visual cortex (from Bregma: posterior, 2.8 mm; lateral, 3.0 mm; ventral, 1.2 mm) of adult mice, anesthetized with 2.5% isoflurane in 100% O_2_, as previously described (Murase et al., 2017; Zold and Hussain Shuler, 2015). Animals received a single dose of Carprofen (5 mg/kg, SQ) for post-surgical analgesia after the return of the righting reflex. Animals were allowed one week to recover from surgery. One day before electrophysiological recordings, animals were habituated for 45 minutes to head restraint. Broad band electrophysiological data was collected from awake head-fixed mice, using RZ5 bioamp processor and RA16PA preamp (Tucker Davis Technologies, TDT). The signal was filtered online for multiunit activity (300 Hz high pass and 5 kHz low pass filtered) and local field potentials (LFP, 300 Hz low pass filter with 60 Hz notch-filter for line noise). Multiunit waveforms were detected using a −4.5 standard deviation threshold then sorted into single units using an automatic Bayesian clustering algorithm in OpenSorter (TDT) utilizing Mahalanobis distance, distance isolation and L-ratio as measures of cluster isolation and quality. Single units were then processed in MATLAB and classified as regular spiking (RS, presumptive excitatory neurons) or fast spiking (FS, presumptive inhibitory neurons) based on waveform slope 0.5 msec after the trough, time between trough and peak, and the ratio of trough to peak height (Niell and Stryker, 2008). Visually evoked potential (VEP) amplitudes were calculated as the amplitude trough to peak of the average of 1 second LFP epochs during visual stimulation, as described (Murase et al., 2017). VEPs and single units were assigned to cortical layer based on LFP waveform shape, and current source density calculated with single site spacing from the laminar array (Guo et al., 2017; Mitzdorf, 1985).

### Visual Stimulation

Visual stimuli were presented using MATLAB (Mathworks) with Psychtoolbox extensions (Brainard, 1997; Pelli, 1997). At the beginning of each recording day animals passively viewed a grey screen of equal luminance equal to grating stimuli (26 cd/m^2^) for 200 seconds (spontaneous). Visually evoked responses were recorded as head-fixed subjects passively viewed 200 × 1s trials of 0.05 cycles per degrees, 100% contrast, square-wave gratings reversing gratings, at various temporal frequencies and orientations. Unless otherwise noted, VEP, single unit and oscillatory activity was recorded during LFVS (200 × 1s trials of 0.05 cycles per degrees, 100% contrast, square-wave gratings reversing at 1 Hz /2 Hz screen refresh). Stimulation frequency is presented as the square wave reversal frequency, followed by the absolute frequency.

### Data Analysis

Spike rates of sorted single units were calculated as the average firing rate during each 1 second epoch (200 trials) and average VEP amplitude was calculated as the peak to trough of the average time-locked LFP using MATLAB. To examine frequency specific changes in oscillatory power, LFP data for evoked and spontaneous stimulations were z-scored, then filtered from 1 to 100 Hz using a sliding frequency window via a bandpass elliptic filter with a span of 3Hz in MATLAB. The analytic signal of these band-passed LFPs was then calculated using a Hilbert transform, and the absolute value was used to calculate power within each frequency band (Fiebelkorn et al., 2018). Power was averaged between trials for each animal, and activity is reported as percent change in power from spontaneous activity during passive viewing of a grey 26 cd/m^2^ grey screen recorded on the first day prior to experimental manipulation (experimental – baseline / baseline × 100). When considering the change in evoked power, averaged percent change in power was calculated during the first 100 ms post stimulus onset, binned oscillatory activity was averaged from this window. Oscillatory bins were defined as: delta 1 - 4 Hz, theta 4 - 8 Hz, alpha 7 - 13 Hz, beta 13 - 30 Hz, and gamma 30 - 100 Hz.

The reliability of incoming visual stimulation to reset the phase of on going oscillations was calculated as Inter-Trial Phase Consistency (ITPC), a time locked measure of oscillatory phase, ranging from 0 when phase is random between trials, to 1 when phase is identical for a given point in time for all trials (Fiebelkorn et al., 2018). To calculate time-locked oscillatory phase, z-scored LFP signals were convolved with Morlet wavelets from 1 to 100 Hz using a 3 Hz window. The cycle width of these wavelets was dependent on the filtered frequency (1 - 10 Hz = 2 cycles, 11 - 14 Hz = 3 cycles, 15 - 20 Hz = 4 cycles, 21 - 100 = 5 cycles). The second half of these complex vectors were normalized, averaged and the absolute value was used to calculate ITPC. To examine the reset of phase during visual stimulation ITPC was averaged during each trial at the onset of visual stimulation. ITPC was binned by oscillatory frequency as follows: delta 1 - 4 Hz, theta 4 - 8 Hz, alpha 7 - 13 Hz, beta 13 - 30 Hz, and gamma 30 - 100 Hz.

### Antibodies

The following antibodies/dilutions were used: rabbit polyclonal anti-c-fos antibody 1:200 (Santa Cruz Biotechnology, Santa Cruz, CA, USA), mouse monoclonal anti-NeuN antibody 1:1000 (Millipore, Bedford, MA, USA), Alexa-488 goat anti-mouse IgG 1:500 and Alexa-546 goat anti-rabbit IgG 1:500 (Molecular Probes, Eugene, OR, USA).

### Immunohistochemistry

Head-fixed subjects passively viewed a grey screen (luminance = 26 cd/m^2^; spontaneous activity) or a high contrast grating reversing at 1 Hz (2 Hz screen refresh), 10 Hz (20 Hz) or 20 Hz (40 Hz) for 200 seconds. 2 hours later subjects were perfused with phosphate buffered saline (PBS) followed by 4% paraformaldehyde (PFA) in PBS. Brains were post-fixed in 4% PFA for 24 hours followed by 30% sucrose for 24 hours, and cryo-protectant solution (0.58 M sucrose, 30% (v/v) ethylene glycol, 3 mM sodium azide, 0.64 M sodium phosphate, pH 7.4) for 24 hours prior to sectioning. Coronal sections (40 μm) were made on a Leica freezing microtome (Model SM 2000R). Sections were blocked with 4% normal goat serum (NGS) in 1X PBS for 1 hour. Antibodies were presented in blocking solution for 18 hours at 4 °C, followed by appropriate secondary antibodies for 6 hr at room temperature.

### Confocal imaging and analysis

Images were acquired on a Zeiss LSM 710 confocal microscope. A maximal intensity projection of the z-stack of binocular primary visual cortex images (from Bregma: AP: −3.0 mm, ML: 2.8 mm, DV: 1.2 mm) was acquired (8 slices at 1.0 μm interval at 512×512 pixels) at 40X (Zeiss Plan-neofluar 40x/1.3 Oil DIC, NA=1.3). c-Fos+ cells were selected by size exclusion (80-300 μm^2^) following NeuN+ colocalization based on fluorescence thresholding (autothreshold+30) and fluorescence intensity (auto threshold + 30). Co-localization of c-fos and NeuN was analyzed in a single Z-section image acquired at 40X, using the JACoP plugin “object-based method” in Fiji. Co-localization was based on centers of mass particle coincidence after ROIs were selected (80-300 μm^2^) using the threshold function (autothreshold+30).

### Statistics

Statistical analysis was completed using JASP (JASP Stats). Repeated measures ANOVA (RANOVA) was used to compare LFP data, arising from 3 time points within the same animal, including VEP, oscillatory power and ITPC, followed by a Bonferroni *post-hoc* when appropriate. We made no assumptions that we recorded from the same single units over multiple days, therefore, a one way ANOVA was employed to compare single unit data from 3 groups, follow by a Tukey *post-hoc* when appropriate. A paired Student’s t-test was used to compare 2 groups of LFP data, and unpaired Student’s t-test was used for 2 groups of single unit data. To compare the change in oscillatory power within subjects we employed a one-sided Student’s t-test. In text values are reported as mean ± standard error, n is reported as the total number of subjects, followed by the total number of recorded single units. Exact p values are reported in the text, except when p < 0.001.

---

**Figure S1. Related to Figure 1C. Supplement Figure 1. HFVS before or after LFVS induces generalized VEP potentiation.** Top: experimental timeline: LFVS (200 presentations of 0.05 cycle per degree 100% contrast gratings 60 degree orientation reversing at 1 Hz/ 2 Hz screen refresh) was preceded by HFVS (200 presentations of 0.05 cycle per degree 100% contrast grating reversing at 10 Hz /20 Hz screen refresh, black) or followed by HFVS (grey). 24 hours later VEPs were evoked with familiar and novel (150 degree orientation) visual stimuli. Similar layer 4 VEP potentiation was observed in both groups in response to familiar or novel visual stimulus (Between subjects RANOVA _(df, 2, 1)_, F=0.01, p = 0.92; n.s. p > 0.05; n = 3 (HFVS preceded LFVS), 11 (HFVS followed LFVS) subjects).

**Figure S2. Related to Figure 1C. Increase in VEP amplitude in response to visual stimuli with novel spatial frequencies and orientation after HFVS.** Significant increase in layer 4 VEP amplitudes 24 hours after HFVS across spatial frequencies (purple) relative to LFVS alone (black) at a novel orientation. (Between groups RANOVA_(df,6,1)_, F= 5.88, p = 0.035; * p < 0.05; n = 9 (HFVS), 6 (LFVS) subjects).

**Figure S3. Related to Figure 1. No VEP potentiation in response to familiar or novel stimulus following stimulation with 5 Hz nor 20 Hz reversing visual stimuli.** A) Experimental timeline: LFVS was followed by 5 Hz reversing stimulus (10 Hz screen refresh, 200 × 1 second presentations of 0.05 cpd, 100% contrast gratings). No change in VEP amplitude in response to familiar or novel visual stimulus in any layer after 24 hours (n = 7 subjects). B) Experimental timeline: LFVS was followed by 20 Hz reversing stimulation (40 Hz screen refresh, 200, 1 second presentations of 0.05 cpd, 100% contrast gratings). No change in VEP amplitude in any cortical layer in response to the familiar or novel visual stimulus after 24 hours (n = 8 subjects).

**Figure S4. Related to Figure 2. 10 Hz visual stimulation increases activity in a large number of neurons.** A) Representative confocal images from layer 4 of V1 from subjects stimulated with 200 seconds of a grey screen (spontaneous, spont, 26 cd/m^2^), or high contrast square wave reversing gratings at 1 Hz (2 Hz screen refresh), 10 Hz (20 Hz) or 20 Hz (40 Hz), stained for c-fos and counter stained with NeuN and DAPI. B) 10 Hz reversing gratings (20 Hz) induce a significant increase in c-fos expression compared to spontaneous and 20 Hz (40 Hz) stimulation (ANOVA _(df, 2, 16)_, F= 6.46, p = 0.004; * = p < 0.05, Bonferroni *post-hoc*; n = 5 subjects (spontaneous), 5 subjects (1 Hz /2 Hz), 5 subjects (10 Hz /20 Hz), 5 subjects (20 Hz /40 Hz). Right: no visually-induced increase in c-fos expression in the non-visual medial prefrontal cortex (mPFC) in same subjects. C) 10 Hz reversing (20 Hz) visual stimulation induces a significant increase the average firing rates of layer 4 RS neurons, compared to 20 Hz (40 Hz) (ANOVA _(df, 2, 65)_, F = 4.49, p = 0.005; * = p < 0.05, Bonferroni *post hoc*; n = 20 units, 11 subjects (10Hz /20 Hz), 17 units, 8 subjects (20Hz /40 Hz), 31 units, 16 subjects (1 Hz /2 Hz).

**Figure S5. Related to Figure 2. Oscillatory power and phase reflect frequency of ongoing visual stimulation.** Left: heat map depicts average oscillatory power from 0 to 100 Hz (3 Hz bins) across cortical layers for first 0.5 seconds of each trial, calculated from the analytic signal binned from LFP by layer. Power is normalized to pre-experimental spontaneous cortical activity. Average power over entire trial, binned within each frequency band (delta: 1-4, theta: 4-8, alpha: 7-13, beta: 13-30, gamma: 30-100). Left; ITPC from 0 to 100 Hz (in 3 Hz bins) for 0.5 seconds of each trial, calculated using Morlet wavelet convolution. Average ITPC over entire trial. A) 5 Hz reversing (10 Hz screen refresh, 200 × 1 second presentations of 0.05 cpd, 100% contrast gratings) visual stimulation significantly decreases oscillatory power in gamma (one-sided t-test, yellow highlight = p < 0.05; n = 7 subjects) and alpha (one-sided t-test, yellow highlight = p < 0.05; n = 7 subjects) while increasing power in theta (one-sided t-test, yellow highlight = p < 0.05; n = 7 subjects) across all cortical layers. B) 5 Hz reversing (10 Hz) visual stimulation increases ITPC in all layers and all oscillatory frequencies above delta (paired t-test. yellow highlight = p < 0.05; n = 7 subjects). C) 20 Hz reversing (40 Hz screen refresh, 200 × 1 second presentations of 0.05 cpd, 100% contrast gratings) visual stimulation significantly increases oscillatory power in beta in layers 2/3 and 4 (One-sided t-test, yellow highlight = p < 0.05; n = 8 subjects), and increases delta power in layer 5/6 (one-sided t-test, yellow highlight = p < 0.05; n = 8 subjects). Theta power is significantly reduced in all cortical layers during 20 Hz reversing (40 Hz) visual stimulation (One-sided t-test, yellow highlight = p < 0.05; n = 8 subjects). D) 20 Hz reversing (40 Hz) visual stimulation increases ITPC, compared to spontaneous, in all layers in beta and gamma frequency bands (paired t-test, yellow highlight = p < 0.05; n = 8 subjects). 20 Hz reversing (40 Hz) visual stimulation decreases theta power in layers 2/3 and layer 4 (paired t-test, yellow highlight = p < 0.05; n = 8 subjects), as well as delta power in layer 2/3 alone (paired t-test, yellow highlight = p < 0.05; n = 8 subjects).

## References

Alain, C., Zhu, K. Da, He, Y., and Ross, B. (2015). Sleep-dependent neuroplastic changes during auditory perceptual learning. Neurobiol. Learn. Mem. 118, 133–142.

Albouy, P., Weiss, A., Baillet, S., and Zatorre, R.J. (2017). Selective Entrainment of Theta Oscillations in the Dorsal Stream Causally Enhances Auditory Working Memory Performance. Neuron 94, 193–206.e5.

Andermann, M.L., Kerlin, A.M., and Reid, C. (2010). Chronic cellular imaging of mouse visual cortex during operant behavior and passive viewing. Front. Cellullar Neurosci. 4, 3.

Aton, S.J., Suresh, A., Broussard, C., and Frank, M.G. (2014). Sleep Promotes Cortical Response Potentiation Following Visual Experience. Sleep 37, 1163–1170.

Bastos, A.M., Vezoli, J., Bosman, C.A., Schoffelen, J.-M., Oostenveld, R., Dowdall, J.R., De Weerd, P., Kennedy, H., and Fries, P. (2015). Visual Areas Exert Feedforward and Feedback Influences through Distinct Frequency Channels. Neuron 85, 390–401.

Bekisz, M., and Wróbel, A. (1999). Coupling of beta and gamma activity in corticothalamic system of cats attending to visual stimuli. Neuroreport 10, 3589–3594.

Beste, C., Wascher, E., Güntürkün, O., and Dinse, H.R. (2011). Improvement and Impairment of Visually Guided Behavior through LTP- and LTD-like Exposure-Based Visual Learning. Curr. Biol. 21, 876–882.

Bortone, D.S., Olsen, S.R., and Scanziani, M. (2014). Translaminar Inhibitory Cells Recruited by Layer 6 Corticothalamic Neurons Suppress Visual Cortex. Neuron 82, 474–485.

Brainard, D.H. (1997). The Psychophysics Toolbox. Spat. Vis. 10, 433–436.

Brickwedde, M., Krüger, M.C., and Dinse, H.R. (2019). Somatosensory alpha oscillations gate perceptual learning efficiency. Nat. Commun. 10, 263.

Camilleri, R., Pavan, A., Ghin, F., Battaglini, L., and Campana, G. (2014). Improvement of uncorrected visual acuity and contrast sensitivity with perceptual learning and transcranial random noise stimulation in individuals with mild myopia. Front. Psychol. 5, 1234.

Cardin, J.A., Carlén, M., Meletis, K., Knoblich, U., Zhang, F., Deisseroth, K., Tsai, L.-H., and Moore, C.I. (2009). Driving fast-spiking cells induces gamma rhythm and controls sensory responses. Nature 459, 663–667.

Chang, M.C., Park, J.M., Pelkey, K.A., Grabenstatter, H.L., Xu, D., Linden, D.J., Sutula, T.P., McBain, C.J., and Worley, P.F. (2010). Narp regulates homeostatic scaling of excitatory synapses on parvalbumin-expressing interneurons. Nat. Neurosci. 13, 1090–1097.

Chen, G., Zhang, Y., Li, X., Zhao, X., Ye, Q., Lin, Y., Tao, H.W., Rasch, M.J., and Zhang, X. (2017). Distinct Inhibitory Circuits Orchestrate Cortical beta and gamma Band Oscillations. Neuron 96, 1403–1418.e6.

Clapp, W.C., Eckert, M.J., Teyler, T.J., and Abraham, W.C. (2006). Rapid visual stimulation induces N-methyl-D-aspartate receptor-dependent sensory long-term potentiation in the rat cortex. Neuroreport 17, 511–515.

Cooke, S.F., and Bear, M.F. (2010). Visual experience induces long-term potentiation in the primary visual cortex. J. Neurosci. 30, 16304–16313.

Cooke, S.F., Komorowski, R.W., Kaplan, E.S., Gavornik, J.P., and Bear, M.F. (2015). Visual recognition memory, manifested as long-term habituation, requires synaptic plasticity in V1. Nat. Neurosci. 18, 262–271.

Crist, R.E., Li, W., and Gilbert, C.D. (2001). Learning to see: experience and attention in primary visual cortex. Nat. Neurosci. 4, 519–525.

Daw, N.W., Fox, K., Sato, H., and Czepita, D. (1992). Critical period for monocular deprivation in the cat visual cortex. J. Neurophysiol. 67, 197–202.

Deveau, J., and Seitz, A.R. (2014). Applying perceptual learning to achieve practical changes in vision. Front. Psychol. 5, 1166.

Durand, S., Iyer, R., Mizuseki, K., de Vries, S., Mihalas, S., and Reid, R.C. (2016). A Comparison of Visual Response Properties in the Lateral Geniculate Nucleus and Primary Visual Cortex of Awake and Anesthetized Mice. J. Neurosci. 36, 12144–12156.

Durkin, J., Suresh, A.K., Colbath, J., Broussard, C., Wu, J., Zochowski, M., and Aton, S.J. (2017). Cortically coordinated NREM thalamocortical oscillations play an essential, instructive role in visual system plasticity. Proc. Natl. Acad. Sci. U. S. A. 114, 10485–10490.

Eaton, N.C., Sheehan, H.M., and Quinlan, E.M. (2016). Optimization of visual training for full recovery from severe amblyopia in adults. Learn. Mem. 23, 99–103.

Espinosa, J.S., and Stryker, M.P. (2012). Development and Plasticity of the Primary Visual Cortex. Neuron 75, 230–249.

Fiebelkorn, I.C., Pinsk, M.A., and Kastner, S. (2018). A Dynamic Interplay within the Frontoparietal Network Underlies Rhythmic Spatial Attention. Neuron 99, 842–853.e8.

Fiorentini, A., and Berardi, N. (1981). Learning in grating waveform discrimination: Specificity for orientation and spatial frequency. Vision Res. 21, 1149–1158.

Frenkel, M.Y., Sawtell, N.B., Diogo, A.C., Yoon, B., Neve, R.L., and Bear, M.F. (2006). Instructive effect of visual experience in mouse visual cortex. Neuron 51, 339–349.

Fuentemilla, L., Marco-Pallarés, J., Münte, T.F., and Grau, C. (2008). Theta EEG oscillatory activity and auditory change detection. Brain Res. 1220, 93–101.

Gavornik, J.P., and Bear, M.F. (2014). Learned spatiotemporal sequence recognition and prediction in primary visual cortex. Nat. Neurosci. 17, 732–737.

Gordon, J.A., and Stryker, M.P. (1996). Experience-dependent plasticity of binocular responses in the primary visual cortex of the mouse. J. Neurosci. 16, 3274–3286.

Gu, Y., Huang, S., Chang, M.C., Worley, P., Kirkwood, A., and Quinlan, E.M. (2013). Obligatory role for the immediate early gene NARP in critical period plasticity. Neuron 79, 335.

Gu, Y., Tran, T., Murase, S., Borrell, A., Kirkwood, A., and Quinlan, E.M. (2016). Neuregulin-Dependent Regulation of Fast-Spiking Interneuron Excitability Controls the Timing of the Critical Period. J. Neurosci. 36, 10285–10295.

Guo, W., Clause, A.R., Barth-Maron, A., and Polley, D.B. (2017). A Corticothalamic Circuit for Dynamic Switching between Feature Detection and Discrimination. Neuron 95, 180–194.e5.

Hanslmayr, S., Axmacher, N., and Inman, C.S. (2019). Modulating Human Memory via Entrainment of Brain Oscillations. Trends Neurosci. 42, 485–499.

He, H.Y., Hodos, W., and Quinlan, E.M. (2006). Visual deprivation reactivates rapid ocular dominance plasticity in adult visual cortex. J. Neurosci. 26, 2951–2955.

Heinze, H.J., Mangun, G.R., Burchert, W., Hinrichs, H., Scholz, M., Münte, T.F., Gös, A., Scherg, M., Johannes, S., Hundeshagen, H., et al. (1994). Combined spatial and temporal imaging of brain activity during visual selective attention in humans. Nature 372, 543–546.

Hillyard, S.A., and Anllo-Vento, L. (1998). Event-related brain potentials in the study of visual selective attention. Proc. Natl. Acad. Sci. U. S. A. 95, 781–787.

Hordacre, B., Ridding, M.C., and Goldsworthy, M.R. (2015). Response variability to non-invasive brain stimulation protocols. Clin. Neurophysiol. 126, 2249–2250.

Howe, W.M., Gritton, H.J., Lusk, N.A., Roberts, E.A., Hetrick, V.L., Berke, J.D., and Sarter, M. (2017). Acetylcholine Release in Prefrontal Cortex Promotes Gamma Oscillations and Theta-Gamma Coupling during Cue Detection. J. Neurosci. 37, 3215–3230.

Huang, S., Gu, Y., Quinlan, E.M., and Kirkwood, A. (2010). A refractory period for rejuvenating GABAergic synaptic transmission and ocular dominance plasticity with dark exposure. J. Neurosci. 30, 16636–16642.

Huang, Y.-Z., Sommer, M., Thickbroom, G., Hamada, M., Pascual-Leonne, A., Paulus, W., Classen, J., Peterchev, A. V., Zangen, A., and Ugawa, Y. (2009). Consensus: New methodologies for brain stimulation. Brain Stimul. 2, 2–13.

Jiang, B., Trevino, M., and Kirkwood, A. (2007). Sequential development of long-term potentiation and depression in different layers of the mouse visual cortex. J. Neurosci. 27, 9648–9652.

Joon Kim, Y., Grabowecky, M., Paller, K.A., Muthu, K., and Suzuki, S. (2007). Attention induces synchronization-based response gain in steady-state visual evoked potentials. Nat. Neurosci. 10, 117–125.

Jutras, M.J., Fries, P., and Buffalo, E.A. (2009). Gamma-band synchronization in the macaque hippocampus and memory formation. J. Neurosci. 29, 12521–12531.

Kaneko, M., Fu, Y., and Stryker, M.P. (2017). Locomotion Induces Stimulus-Specific Response Enhancement in Adult Visual Cortex. J. Neurosci. 37, 3532–3543.

Kaplan, E.S., Cooke, S.F., Komorowski, R.W., Chubykin, A.A., Thomazeau, A., Khibnik, L.A., Gavornik, J.P., and Bear, M.F. (2016). Contrasting roles for parvalbumin-expressing inhibitory neurons in two forms of adult visual cortical plasticity. Elife 5.

Karawani, H., Bitan, T., Attias, J., and Banai, K. (2016). Auditory Perceptual Learning in Adults with and without Age-Related Hearing Loss. Front. Psychol. 6, 2066.

Karni, A., and Sagi, D. (1993). The time course of learning a visual skill. Nature 365, 250–252.

Karni, A., Tanne, D., Rubenstein, B.S., Askenasy, J.J., and Sagi, D. (1994). Dependence on REM sleep of overnight improvement of a perceptual skill. Science 265, 679–682.

van Kerkoerle, T., Self, M.W., Dagnino, B., Gariel-Mathis, M.-A., Poort, J., van der Togt, C., and Roelfsema, P.R. (2014). Alpha and gamma oscillations characterize feedback and feedforward processing in monkey visual cortex. Proc. Natl. Acad. Sci. U. S. A. 111, 14332–14341.

Kissinger, S.T., Pak, A., Tang, Y., Masmanidis, S.C., and Chubykin, A.A. (2018). Oscillatory Encoding of Visual Stimulus Familiarity. J. Neurosci. 38, 6223–6240.

Levi, D.M., and Li, R.W. (2009). Perceptual learning as a potential treatment for amblyopia: A mini-review. Vision Res. 49, 2535–2549.

Lisman, J. (2010). Working Memory: The Importance of Theta and Gamma Oscillations. Curr. Biol. 20, R490–R492.

Marzoll, A., Saygi, T., and Dinse, H.R. (2018). The effect of LTP- and LTD-like visual stimulation on modulation of human orientation discrimination. Sci. Rep. 8, 16156.

Mitzdorf, U. (1985). Current source-density method and application in cat cerebral cortex: investigation of evoked potentials and EEG phenomena. Physiol. Rev. 65, 37–100.

Montey, K.L., and Quinlan, E.M. (2011). Recovery from chronic monocular deprivation following reactivation of thalamocortical plasticity by dark exposure. Nat.Commun. 2, 317.

Montgomery, S.M., and Buzsáki, G. (2007). Gamma oscillations dynamically couple hippocampal CA3 and CA1 regions during memory task performance. Proc. Natl. Acad. Sci. U. S. A. 104, 14495–14500.

Moretti, D.V., Fracassi, C., Pievani, M., Geroldi, C., Binetti, G., Zanetti, O., Sosta, K., Rossini, P.M., and Frisoni, G.B. (2009). Increase of theta/gamma ratio is associated with memory impairment. Clin. Neurophysiol. 120, 295–303.

Morgan, S.T., Hansen, J.C., and Hillyard, S.A. (1996). Selective attention to stimulus location modulates the steady-state visual evoked potential. Proc. Natl. Acad. Sci. U. S. A. 93, 4770–4774.

Murase, S., Lantz, C.L., and Quinlan, E.M. (2017). Light reintroduction after dark exposure reactivates plasticity in adults via perisynaptic activation of MMP-9. Elife 6.

Niell, C.M., and Stryker, M.P. (2008). Highly selective receptive fields in mouse visual cortex. J. Neurosci. 28, 7520–7536.

O’Brien, R.J., Xu, D., Petralia, R.S., Steward, O., Huganir, R.L., and Worley, P. (1999). Synaptic clustering of AMPA receptors by the extracellular immediate-early gene product Narp. Neuron 23, 309–323.

Olsen, S.R., Bortone, D.S., Adesnik, H., and Scanziani, M. (2012). Gain control by layer six in cortical circuits of vision. Nature 483, 47–52.

Park, H., Lee, D.S., Kang, E., Kang, H., Hahm, J., Kim, J.S., Chung, C.K., Jiang, H., Gross, J., and Jensen, O. (2016). Formation of visual memories controlled by gamma power phase-locked to alpha oscillations. Sci. Rep. 6, 28092.

Pelli, D.G. (1997). The VideoToolbox software for visual psychophysics: transforming numbers into movies. Spat. Vis. 10, 437–442.

Poggio, T., Fahle, M., Edelman, S., Askenasy, J., and Sagi, D. (1992). Fast perceptual learning in visual hyperacuity. Science (80-.). 256, 1018–1021.

Porciatti, V., Pizzorusso, T., and Maffei, L. (1999). The visual physiology of the wild type mouse determined with pattern VEPs. Vision Res. 39, 3071–3081.

Ren, Z., Zhou, J., Yao, Z., Wang, Z., Yuan, N., Xu, G., Wang, X., Zhang, B., Hess, R.F., and Zhou, Y. (2016). Neuronal basis of perceptual learning in striate cortex. Sci. Rep. 6, 24769.

Sagi, D. (2011). Perceptual learning in Vision Research. Vision Res. 51, 1552–1566.

Sale, A., and Berardi, N. (2015). Active training for amblyopia in adult rodents. Front. Behav. Neurosci. 9, 281.

Schoups, A.A., Vogels, R., and Orban, G.A. (1995). Human perceptual learning in identifying the oblique orientation: retinotopy, orientation specificity and monocularity. J. Physiol. 483, 797–810.

Seitz, A.R. (2017). Perceptual learning. Curr. Biol. 27, R631–R636.

Sohal, V.S., Zhang, F., Yizhar, O., and Deisseroth, K. (2009). Parvalbumin neurons and gamma rhythms enhance cortical circuit performance. Nature 459, 698–702.

Spaak, E., Bonnefond, M., Maier, A., Leopold, D.A., and Jensen, O. (2012). Layer-Specific Entrainment of Gamma-Band Neural Activity by the Alpha Rhythm in Monkey Visual Cortex. Curr. Biol. 22, 2313–2318.

Stothart, G., and Kazanina, N. (2013). Oscillatory characteristics of the visual mismatch negativity: what evoked potentials aren’t telling us. Front. Hum. Neurosci. 7, 426.

Taaseh, N., Yaron, A., and Nelken, I. (2011). Stimulus-Specific Adaptation and Deviance Detection in the Rat Auditory Cortex. PLoS One 6, e23369.

Tanimoto, N., Sothilingam, V., Kondo, M., Biel, M., Humphries, P., and Seeliger, M.W. (2015). Electroretinographic assessment of rod- and cone-mediated bipolar cell pathways using flicker stimuli in mice. Sci. Rep. 5, 10731.

Teyler, T.J., Hamm, J.P., Clapp, W.C., Johnson, B.W., Corballis, M.C., and Kirk, I.J. (2005). Long-term potentiation of human visual evoked responses. Eur. J. Neurosci. 21, 2045–2050.

Todd, G., Rogasch, N.C., Flavel, S.C., and Ridding, M.C. (2009). Voluntary movement and repetitive transcranial magnetic stimulation over human motor cortex. J. Appl. Physiol. 106, 1593–1603.

Tsapa, D., Ahmadlou, M., and Heimel, J.A. (2019). Long-term enhancement of visual responses by repeated transcranial electrical stimulation of the mouse visual cortex. Brain Stimul.

Voloh, B., and Womelsdorf, T. (2016). A Role of Phase-Resetting in Coordinating Large Scale Neural Networks During Attention and Goal-Directed Behavior. Front. Syst. Neurosci. 10, 18.

Williams, J.. (2001). Frequency-specific effects of flicker on recognition memory. Neuroscience 104, 283–286.

Williams, J., Ramaswamy, D., and Oulhaj, A. (2006). 10 Hz flicker improves recognition memory in older people. BMC Neurosci. 7, 21.

Xu, D., Hopf, C., Reddy, R., Cho, R.W., Guo, L., Lanahan, A., Petralia, R.S., Wenthold, R.J., O’Brien, R.J., and Worley, P. (2003). Narp and NP1 Form Heterocomplexes that Function in Developmental and Activity-Dependent Synaptic Plasticity. Neuron 39, 513–528.

Zhang, W., and Luck, S.J. (2009). Feature-based attention modulates feedforward visual processing. Nat. Neurosci. 12, 24–25.

Zold, C.L., and Hussain Shuler, M.G. (2015). Theta Oscillations in Visual Cortex Emerge with Experience to Convey Expected Reward Time and Experienced Reward Rate. J. Neurosci. 35, 9603–9614.

