## Supplemental Figures and Legends for "Regulation of expression site and generalizability of experience-dependent plasticity in visual cortex"

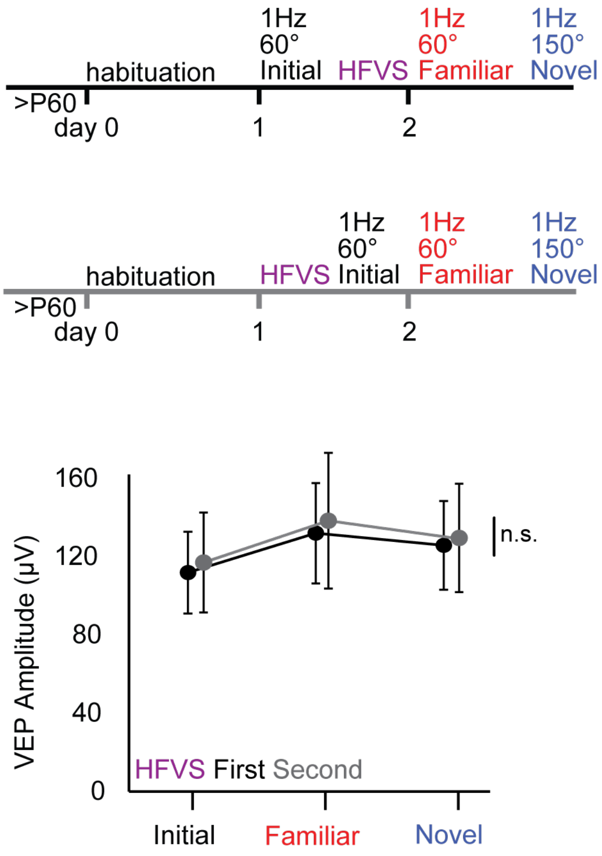

1Hz
60°
Initial
HFVS
1Hz
60°
Familiar
1Hz
150°
Novel
habituation
>P60
day 0
1
2
habituation
HFVS
1Hz
60°
Initial
1Hz
60°
Familiar
1Hz
150°
Novel
>P60
day 0
1
2
VEP Amplitude (μV)
160
120
80
40
0
n.s.
HFVS First Second
Initial
Familiar
Novel


**Figure S1.**

**Figure S1. Related to Figure 1C. Supplement Figure 1. HFVS before or after LFVS induces generalized VEP potentiation.** Top: experimental timeline: LFVS (200 presentations of 0.05 cycle per degree 100% contrast gratings 60 degree orientation reversing at 1 Hz/ 2 Hz screen refresh) was preceded by HFVS (200 presentations of 0.05 cycle per degree 100% contrast grating reversing at 10 Hz /20 Hz screen refresh, black) or followed by HFVS (grey). 24 hours later VEPs were evoked with familiar and novel (150 degree orientation) visual stimuli. Similar layer 4 VEP potentiation was observed in both groups in response to familiar or novel visual stimulus (Between subjects RANOVA $_{(df, 2, 1)}$, $F=0.01$, $p = 0.92$; n.s. $p > 0.05$; $n = 3$ (HFVS preceded LFVS), 11 (HFVS followed LFVS) subjects).

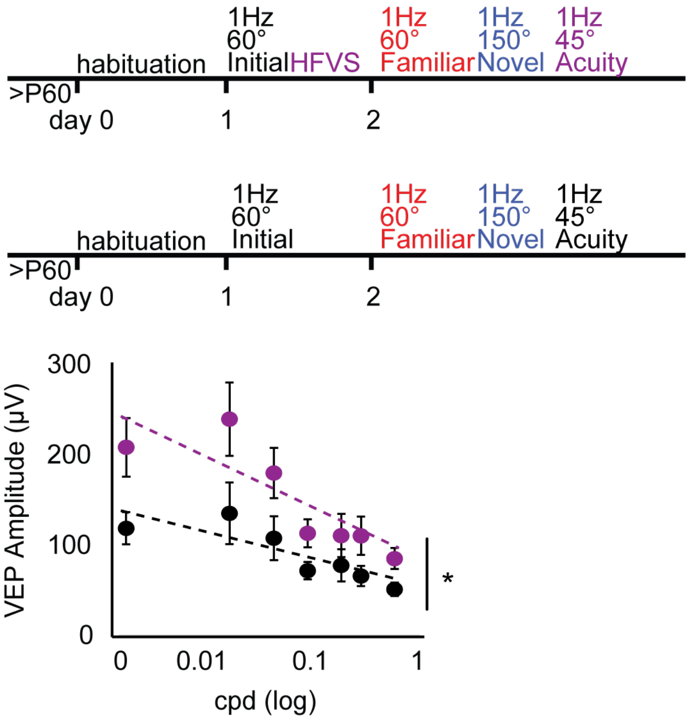
1Hz
60°
Initial
HFVS
1Hz
60°
Familiar
1Hz
150°
Novel
1Hz
45°
Acuity
habituation
>P60
day 0
1
2
1Hz
60°
Initial
1Hz
60°
Familiar
1Hz
150°
Novel
1Hz
45°
Acuity
habituation
>P60
day 0
1
2
VEP Amplitude (µV)
300
200
100
0
0
0.01
0.1
1
cpd (log)
*

**Figure S2.**

**Figure S2. Related to Figure 1C. Increase in VEP amplitude in response to visual stimuli with novel spatial frequencies and orientation after HFVS.** Significant increase in layer 4 VEP amplitudes 24 hours after HFVS across spatial frequencies (purple) relative to LFVS alone (black) at a novel orientation. (Between groups $RANOVA_{(df,6,1)}$, $F = 5.88$, $p = 0.035$; * $p < 0.05$; n = 9 (HFVS), 6 (LFVS) subjects).

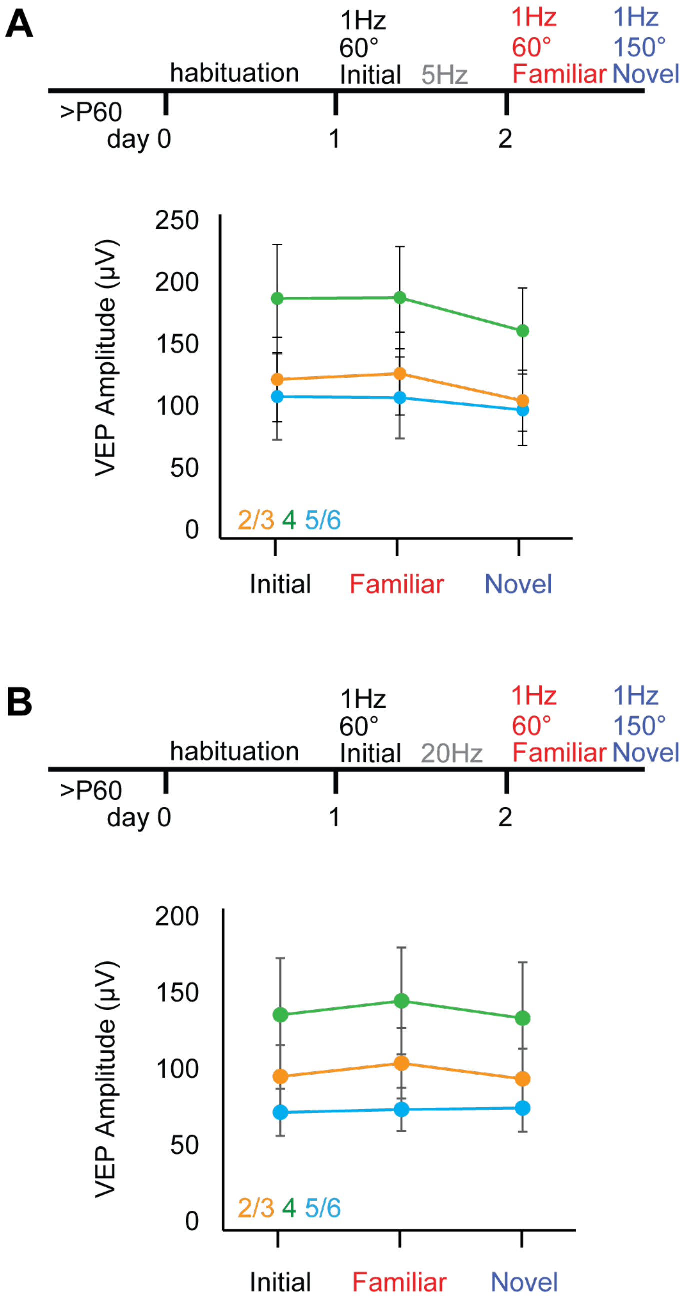
A
habituation
1Hz
60°
Initial
5Hz
1Hz
60°
Familiar
1Hz
150°
Novel
>P60
day 0
1
2
VEP Amplitude (µV)
250
200
150
100
50
0
2/3 4 5/6
Initial
Familiar
Novel
B
habituation
1Hz
60°
Initial
20Hz
1Hz
60°
Familiar
1Hz
150°
Novel
>P60
day 0
1
2
VEP Amplitude (µV)
200
150
100
50
0
2/3 4 5/6
Initial
Familiar
Novel

**Figure S3.**

**Figure S3. Related to Figure 1. No VEP potentiation in response to familiar or novel stimulus following stimulation with 5 Hz nor 20 Hz reversing visual stimuli.** A) Experimental timeline: LFVS was followed by 5 Hz reversing stimulus (10 Hz screen refresh, 200 x 1 second presentations of 0.05 cpd, 100% contrast gratings). No change in VEP amplitude in response to familiar or novel visual stimulus in any layer after 24 hours (n = 7 subjects). B) Experimental timeline: LFVS was followed by 20 Hz reversing stimulation (40 Hz screen refresh, 200, 1 second presentations of 0.05 cpd, 100% contrast gratings). No change in VEP amplitude in any cortical layer in response to the familiar or novel visual stimulus after 24 hours (n = 8 subjects).

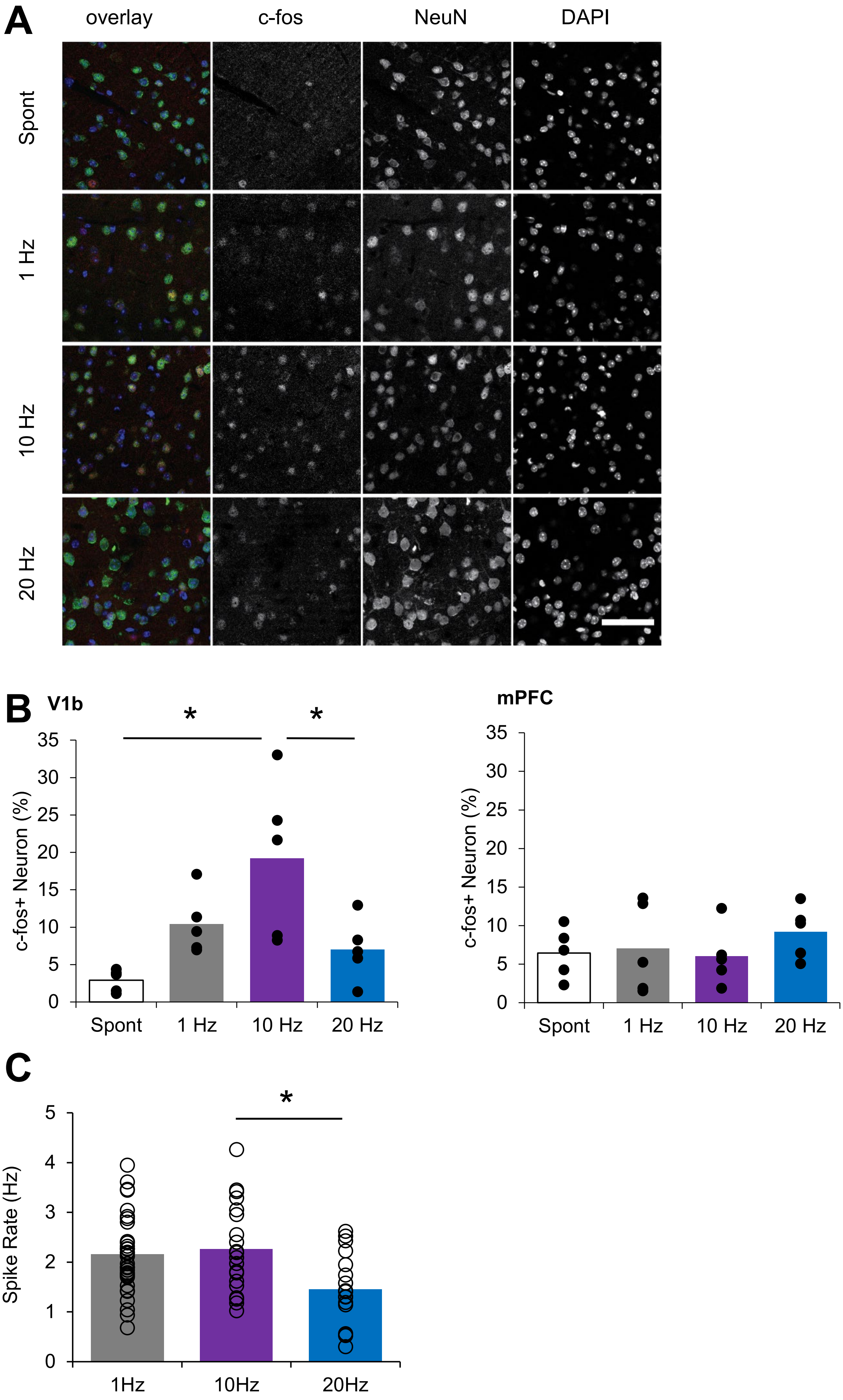
A
overlay
c-fos
NeuN
DAPI
Spont
1 Hz
10 Hz
20 Hz
B
V1b
*
*
c-fos+ Neuron (%)
0
5
10
15
20
25
30
35
Spont
1 Hz
10 Hz
20 Hz
mPFC
c-fos+ Neuron (%)
0
5
10
15
20
25
30
35
Spont
1 Hz
10 Hz
20 Hz
C
*
Spike Rate (Hz)
0
1
2
3
4
5
1Hz
10Hz
20Hz

**Figure S4.**

**Figure S4. Related to Figure 2. 10 Hz visual stimulation increases activity in a large number of neurons.** A) Representative confocal images from layer 4 of V1 from subjects stimulated with 200 seconds of a grey screen (spontaneous, spont, 26 $cd/m^2$), or high contrast square wave reversing gratings at 1 Hz (2 Hz screen refresh), 10 Hz (20 Hz) or 20 Hz (40 Hz), stained for c-fos and counter stained with NeuN and DAPI. B) 10 Hz reversing gratings (20 Hz) induce a significant increase in c-fos expression compared to spontaneous and 20 Hz (40 Hz) stimulation (ANOVA $_{(df, 2, 16)}$, F= 6.46, p = 0.004; * = p < 0.05, Bonferroni *post-hoc*; n = 5 subjects (spontaneous), 5 subjects (1 Hz /2 Hz), 5 subjects (10 Hz /20 Hz), 5 subjects (20 Hz /40 Hz). Right: no visually-induced increase in c-fos expression in the non-visual medial prefrontal cortex (mPFC) in same subjects. C) 10 Hz reversing (20 Hz) visual stimulation induces a significant increase the average firing rates of layer 4 RS neurons, compared to 20 Hz (40 Hz) (ANOVA $_{(df, 2, 65)}$, F = 4.49, p = 0.005; * = p < 0.05, Bonferroni *post hoc*; n = 20 units, 11 subjects (10Hz /20 Hz), 17 units, 8 subjects (20Hz /40 Hz), 31 units, 16 subjects (1 Hz /2 Hz).

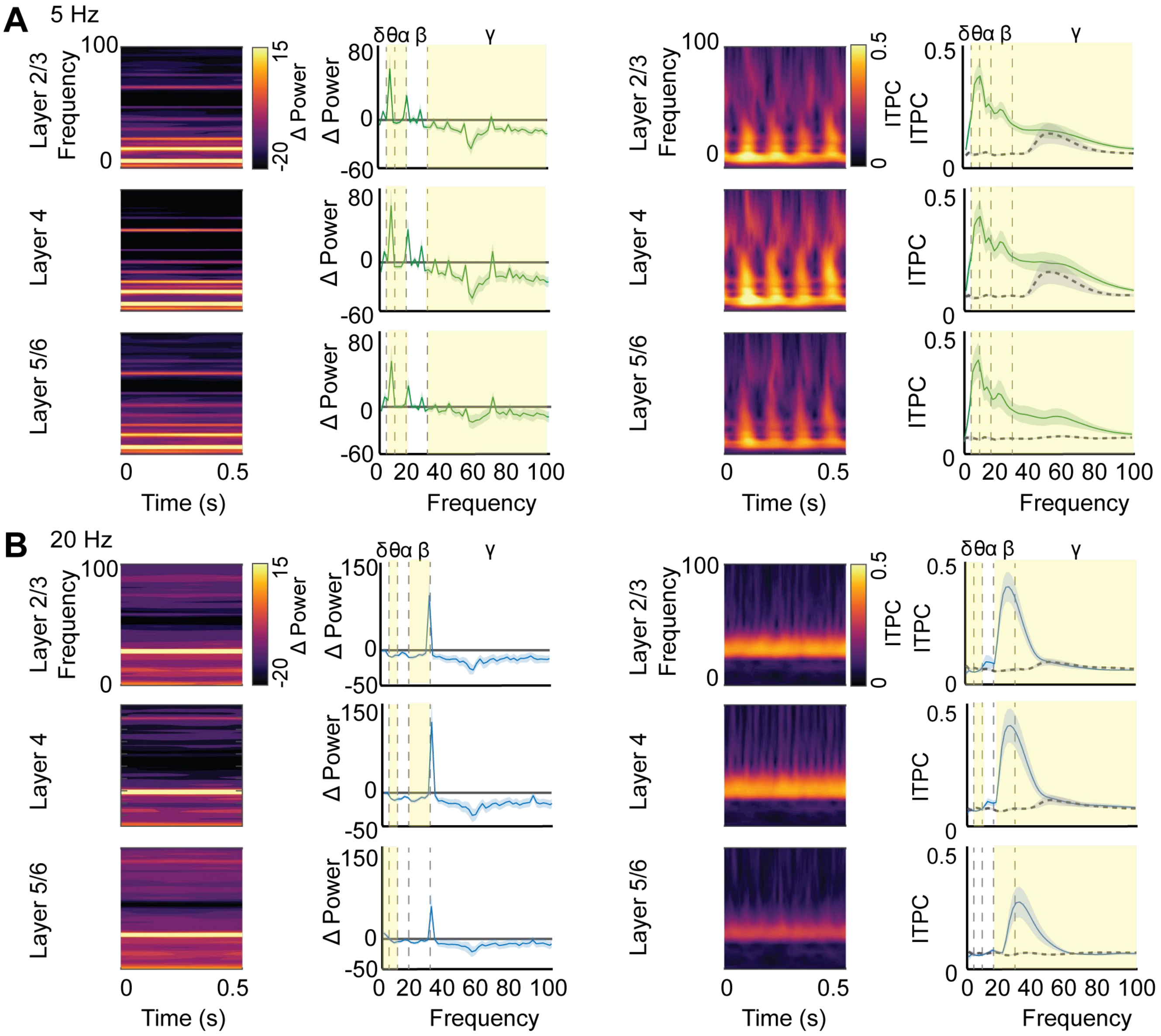

A
5 Hz
Layer 2/3
Layer 4
Layer 5/6
Frequency
Time (s)
Δ Power
ITPC
δ θ α β
γ
B
20 Hz


**Figure S5.**

**Figure S5. Related to Figure 2. Oscillatory power and phase reflect frequency of ongoing visual stimulation.** Left: heat map depicts average oscillatory power from 0 to 100 Hz (3 Hz bins) across cortical layers for first 0.5 seconds of each trial, calculated from the analytic signal binned from LFP by layer. Power is normalized to pre-experimental spontaneous cortical activity. Average power over entire trial, binned within each frequency band (delta: 1-4, theta: 4-8, alpha: 7-13, beta: 13-30, gamma: 30-100). Left; ITPC from 0 to 100 Hz (in 3 Hz bins) for 0.5 seconds of each trial, calculated using Morlet wavelet convolution. Average ITPC over entire trial. A) 5 Hz reversing (10 Hz screen refresh, 200 x 1 second presentations of 0.05 cpd, 100% contrast gratings) visual stimulation significantly decreases oscillatory power in gamma (one-sided t-test, yellow highlight = $p < 0.05$; $n = 7$ subjects) and alpha (one-sided t-test, yellow highlight = $p < 0.05$; $n = 7$ subjects) while increasing power in theta (one-sided t-test, yellow highlight = $p < 0.05$; $n = 7$ subjects) across all cortical layers. B) 5 Hz reversing (10 Hz) visual stimulation increases ITPC in all layers and all oscillatory frequencies above delta (paired t-test. yellow highlight = $p < 0.05$; $n = 7$ subjects). C) 20 Hz reversing (40 Hz screen refresh, 200 x 1 second presentations of 0.05 cpd, 100% contrast gratings) visual stimulation significantly increases oscillatory power in beta in layers 2/3 and 4 (One-sided t-test, yellow highlight = $p < 0.05$; $n = 8$ subjects), and increases delta power in layer 5/6 (one-sided t-test, yellow highlight = $p < 0.05$; $n = 8$ subjects). Theta power is significantly reduced in all cortical layers during 20 Hz reversing (40 Hz) visual stimulation (One-sided t-test, yellow highlight = $p < 0.05$; $n = 8$ subjects). D) 20 Hz reversing (40 Hz) visual stimulation increases ITPC, compared to spontaneous, in all layers in beta and gamma frequency bands (paired t-test, yellow highlight = $p < 0.05$; $n = 8$ subjects). 20 Hz reversing (40 Hz) visual stimulation decreases theta power in layers 2/3 and layer 4 (paired t-test, yellow highlight = $p < 0.05$; $n = 8$ subjects), as well as delta power in layer 2/3 alone (paired t-test, yellow highlight = $p < 0.05$; $n = 8$ subjects).
